# Proline transporters balance the salicylic acid-mediated trade-off between regeneration and immunity in plants

**DOI:** 10.1101/2025.11.20.689487

**Authors:** Dawei Xu, Zeinu Mussa Belew, Nathália Cássia Ferreira Dias, Li Wang, Andy Zhang, Yun-Fan Stephanie Chen, Carter James Newton, Feng Kong, Yaochao Zheng, Yao Yao, Marin Talbot Brewer, Paulo José Pereira Lima Teixeira, Hussam Hassan Nour Eldin Auis, Deyang Xu, Li Yang

## Abstract

A robust immune response and regenerative capacity are both essential for survival after injury. In plants, salicylic acid (SA) is essential for activating immunity but simultaneously suppresses regenerative capacity. The mechanisms coordinating the trade-off between immunity and regeneration remain poorly understood. Here, we identify proline transporters as key regulators of this balance. Mutations in two wound-induced proline transporters, ProT2 and ProT3, rescued exogenous proline-induced suppression of de novo root regeneration (DNRR) and enhanced DNRR. ProTs are required for SA–mediated suppression of regeneration, without affecting SA-dependent defense responses. Mechanistically, a ProT3–CPK1 complex modulates the dynamics of wound-induced reactive oxygen species (ROS), sustaining a ROS level that restricts DNRR after wounding. Notably, pharmacological inhibition of proline transport rescued SA-mediated suppression of regeneration and enhanced regeneration across multiple plant species. These findings establish proline transporter as a regulatory hub integrating stress-induced proline metabolism, SA signaling, and ROS homeostasis to balance immunity and regeneration, and highlight chemical inhibition of proline transport as a strategy to improve crop regeneration under biotic stresses without compromising disease resistance.

## Introduction

Regeneration and immunity are two fundamental processes that enable plants to survive wounding and other forms of tissue damage caused by pathogen invasion, herbivore feeding, and mechanical injuries under extreme weather conditions. Upon damage, plants initiate wound healing and, in some cases, produce new organs to restore function ^1^. During de novo root regeneration (DNRR), detached leaves or stem cuttings undergo wound healing and callus formation, leading to adventitious root development, which are guided exclusively by endogenous hormone signals ^2^. At the same time, multiple layers of immune responses are activated locally or systemically in response to wound at a time scale from seconds to days to constrain microbial entry and multiplication ^3, 4^. These responses may include the activation of reactive oxygen species (ROS) bursts, reprogramming of phytohormone signaling, strengthening of cell wall and secretion of antimicrobial compounds ^5^. Although these dual processes—defense and regeneration—are indispensable for plant survival in natural ecosystems, regeneration is often disrupted under high pressure of biotic stresses such as the presence of pathogenic microbes ^6^.

Recent advancements have begun to unveil the molecular mechanisms through which immune activation inhibits the regenerative potential in plants ^7^. For instance, activation of salicylic acid (SA) signaling has been repeatedly shown to impair plant regeneration. SA accumulation after wounding inhibits callus formation in Arabidopsis and maize tissue culture ^8^. Similarly, DNRR from Arabidopsis leaf explants is strongly reduced by exogenous SA treatment or in mutants with constitutively elevated SA levels ^9^. Inside of a callus, SA relays hypoxia stress signal to suppress shoot organogenesis ^10^. Beyond the role of SA, microbe-associated molecular patterns (MAMPs) can also block DNRR; the bacterial elicitor flg22, a 22 amino acids peptide derived from flagellin, prevents adventitious root initiation from wounded tissues ^11^. These examples illustrate that immune signaling—while vital for pathogen defense—can restrict regenerative competence. Thus, fine-tuning the balance between immunity and regeneration is critical for plant survival under combined abiotic and biotic stresses.

Wounding often induces multiple stresses simultaneously ^3, 12^. Proline is a key metabolite positioned at the interface of multiple stress responses ^13, 14^. Under drought, salinity, or oxidative stress, proline accumulates rapidly and may function as an osmoprotectant, stabilizer of proteins and membranes, and regulator of cellular redox homeostasis ^15, 16, 17, 18, 19, 20, 21^. Its levels are tightly controlled by metabolic enzymes such as P5CS (pyrroline-5-carboxylate synthetase) for synthesis and ProDH (proline dehydrogenase) for degradation, as well as by dedicated transporters ^22, 23^. The spatial and temporal distribution of proline in plant tissues is tightly regulated by proline transporters (ProTs), such as the Arabidopsis ProT family (AtProT1, AtProT2, and AtProT3) ^24, 25, 26^. Arabidopsis proline transporter family mediates the uptake of proline as well as γ-aminobutyric acid (GABA), and quaternary ammonium compounds in yeast ^25, 27^. ProTs and other proline transporters such as AMINO ACID PERMEASE1 and Lysine Histidine Transporter 1 (LHT1), link proline metabolism to plant growth, immunity, and stress adaptation ^26, 28^. Recent work shows that MAMP perception triggers LHT1-dependent uptake of proline from the leaf apoplast, and this targeted depletion reinforces pattern-triggered immunity and underscores nutrient restriction in apoplast as a key component of plant defense ^26^.

SA and proline often act synergistically to alleviate plant stress responses. SA promotes proline biosynthesis and catabolism under conditions such as drought, salinity, and heat^15, 19, 29, 30, 31, 32^. Exogenous co-application of SA and proline has been shown to yield additive or synergistic effects in multiple crops to counteract stress-induced defects ^33, 34, 35, 36, 37^. In regeneration contexts, proline displays multifaceted roles. Exogenous proline enhances callus induction and regeneration in *Brassica napus*, banana, and rice ^38, 39, 40^, but in some cases, it reduces somatic embryo numbers (peanut) or inhibits cell growth (saltgrass, *Distichlis spicata*) during regeneration ^41, 42, 43^. These findings suggest that proline influences cellular competence for regeneration in a species-, cell type- and dose-dependent manner, although the underlying mechanism is largely unknown.

In this study, we demonstrate that the Arabidopsis proline transporters ProT2 and ProT3 play key roles in maintaining the balance between SA-mediated defense and the plant’s ability to regenerate after damage. We show that exogenous proline suppresses DNRR in Arabidopsis, correlating with repression of the root founder cell marker *WOX11*. Mutants of *prot2* and *prot3* exhibit enhanced regenerative capacity, whereas overexpression of ProT3 impairs wound-induced adventitious root formation. We identify a ProT3–CPK1 protein complex that restricts regeneration downstream of SA signaling, likely by maintaining elevated wound-induced ROS. Importantly, this complex is dispensable for SA-induced defense gene activation, thereby uncoupling SA-mediated suppression of regeneration and activation of defense. Pharmacological inhibition of proline transporter activity using the small molecule LQFM215 rescues SA-induced regeneration defects and promotes organogenesis in Arabidopsis, tomato, and other species. These findings offer a strategy to enhance crop regenerative capacity without compromising immunity.

## Results

### Members of proline transporter family have distinct roles in regeneration

Our previous work showed that wound-induced SA suppressed DNRR, a process in which roots regenerate from wound sites without exogenous hormones, in Arabidopsis ^9^. Since SA and proline act synergistically in response to various abiotic stresses, we sought to investigate the role of proline in DNRR. We first treated leaf explants with various concentration of L-proline and found that L-proline reduced DNRR at 10 µM and completely blocked adventitious root formation at 10mM (Fig. 1a). Being consistent with the reduced DNRR, L-proline also inhibited the expression of *WOX11*, a marker of cell fate transition from procambium cell to root founder cell during DNRR, at cutting sites (Fig. 1b) ^44^. These results suggest that endogenous proline may negatively influence wound-induced regeneration. The analysis of genes that are induced after leaf cutting revealed that the expression of *ProT3* was rapidly induced after cutting, resembling the expression pattern of some SA-responsive genes (Extended Data Fig. 1a) ^9, 45^. To investigate ProT3’s potential function in regeneration, we first analyzed the DNRR phenotype of Arabidopsis *prot3-1* and *prot3-2* mutants. Both *prot3-1* and *prot3-2* alleles exhibited enhanced adventitious root formation compared to Col-0 (Fig. 1c, d). This enhanced regenerative capacity was restored to wild type level by a genomic *ProT3* construct containing the *ProT3* coding region and 1919 bp upstream, confirming that the phenotype results from a recessive loss-of-function mutation (Extended Data Fig. 1b). We also examined wound-induced callus (WIC), another form of regeneration without exogenous hormone application. In this system, the *prot3-2* mutant developed enlarged calli at wound sites (Fig. 1e, f). Furthermore, two independent lines overexpressing ProT3-YFP under the ubiquitin-10 promoter displayed significantly reduced DNRR (Fig. 1g). Similar to the application of exogenous proline, *ProT3* overexpression also suppressed wound-induced *WOX11*, but not its expression in un-wounded tissues, indicating that ProT3 negatively regulates root founder cell establishment during DNRR (Fig. 1h). We also investigated the DNRR phenotype in mutants of other proline transporter family members: *prot2* mutant enhanced DNRR similar to *prot3*, whereas *prot1* mutant had no effect, suggesting a functional divergence within the ProT family (Extended Data Fig. 1d,e).

**Fig. 1.**
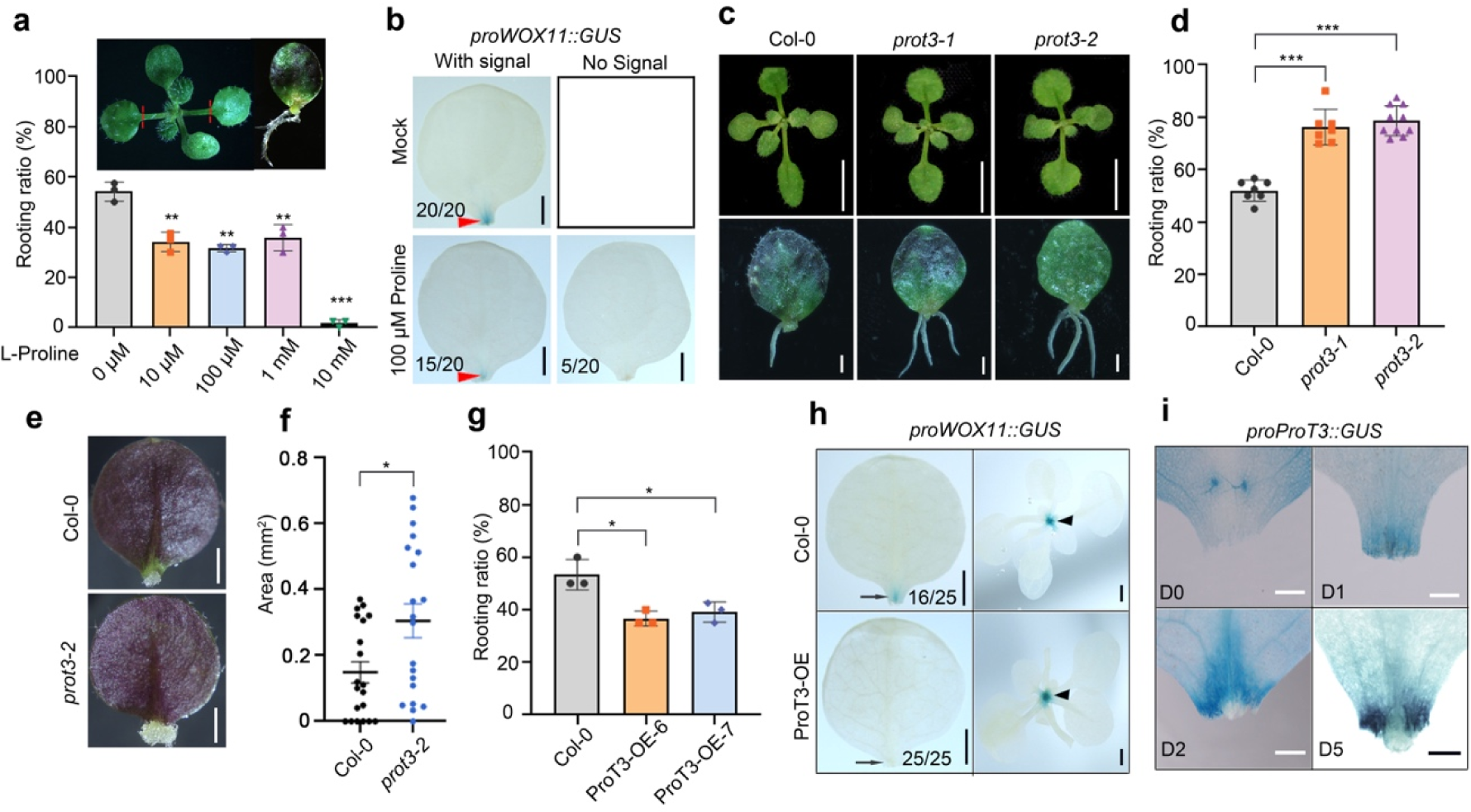
ProT3 is induced by wounding and negatively regulates DNRR. a, Ratio of adventitious root formation on Col-0 leaf explants treated with increasing concentrations of L-proline (0, 10, 100 μM, 1, and 10 mM). (Each dot represents one biological replicate with 40 explants. n=3). The insert shows an Arabidopsis seedling (left) and an explant with adventitious root (right). Red dashed line indicates cutting position. b, Representative images of *proWOX11::GUS* staining in leaf explants treated with 100 μM L-proline at 1 day after cutting (1DAC). scale bar: 1 mm. Red arrowheads indicate the cutting sites. c, Morphology of 11-day-old plants (top) and DNRR phenotypes (bottom) of Col-0, *prot3-1* and *prot3-2*; scale bars, 0.5 cm (top) and 1 mm (bottom). d, Rooting ratio of Col-0, *prot3-1* and *prot3-2* explants cultured for 10 d (n > 7. 40 explants for each repeat). e, f, *prot3* mutation enhanced wound-induced callus (WIC) (e) and the statistical analysis of callus area (f). n = 20 petioles per genotype, scale bar, 1 mm. g. Rooting ratio in Col-0, ProT3-OE-6, and ProT3-OE-7 leaf explants. (n = 3 biological replicates, 40 explants each). h, *proWOX11::GUS* expression in Col-0 and ProT3-OE-7 leaf explants at 1 DAC, scale bar, 1 mm. Arrows and arrowheads indicate GUS activity at cutting sites and shoot-hypocotyle junction, respectively. i, *proProT3::GUS* expression in leaf explants at D0, 1DAC, 2DAC and 5DAC, scale bar, 1 mm. Data are presented as mean ± s.d.; n denotes the number of biological replicates Statistical significance was determined by two-tailed Student’s t-test; *P < 0.05, **P <0.01, ***P < 0.001; ns, not significant.

To further elucidate the spatial and temporal regulation of proline transporters during DNRR, we generated promoter–GUS reporter lines for *ProT1*, *ProT2*, and *ProT3*. Immediately after cutting (day 0, D0), GUS activity driven by the *ProT3* promoter was detected in leaf blades but absent at cutting sites (Fig. 1i). At 1 day after cutting (1 DAC, D1), *ProT3* was strongly induced at the wound site (Fig. 1i). With the development of callus tissue, *proProT3::GUS* activity was excluded from callus, with activity restricted to the peripheral zone (Fig. 1i). Similarly, *proProT2::GUS* was strongly induced at cutting sites at 1DAC (Extended Data Fig. 1f). In contrast, *proProT1::GUS* expression was confined to vascular tissues and was not wound-inducible at 1 DAC (Extended Data Fig. 1g). These patterns are consistent with the roles of *ProT3* and *ProT2*, but not *ProT1*, as negative regulators of wound-induced regeneration. Collectively, these findings highlight the divergent roles of *ProT* family members in DNRR, likely due to their distinct expression dynamics.

### ProT3 suppresses DNRR through its proline transport activity

To test the role of ProT2 and ProT3 in proline-mediated inhibition of DNRR, we first validated proline transport activity of ProTs using *Xenopus laevis* oocyte expression system. Oocytes expressing ProT2 or ProT3 can import ^3^H-proline into the cells from external buffer at pH5.0 but not at pH7.4, compared to the cells expressing GFP (***P < 0.001, n = 10) (Fig. 2a, b). Moreover, ProT3 showed higher ^3^H-proline uptake than ProT2 in 30 minutes import assay (Fig. 2a). Electrophysiological recordings revealed robust proline-induced inward currents in ProT2- and ProT3-expressing oocytes at pH5.0, which were absent in GFP controls (Fig. 2c). These suggested that ProTs-mediated import of proline is driven by proton gradient across plasma membrane. When treated with L-proline at concentrations of 10 µM and 100 µM that inhibited DNRR in Col-0, *prot3* mutants were insensitive to such treatment, indicating that the mutant is defective in importing extracellular proline (Fig. 2d). At 10 mM, L-proline treatment inhibited DNRR in *prot3* mutants, probably due to toxicity or the function of other low-affinity proline transporters (Fig. 2d). In contrast, the suppression of DNRR by mannitol was comparable in both Col-0 and *prot3* mutants (Fig. 2e), suggesting that ProT3’s role is specifically required for proline-induced inhibition of DNRR rather than a reduced sensitivity to osmolality. Given that ProTs are localized to the plasma membrane ^25^, these findings suggest that ProT3-mediated proline import into cells at the wound site, where it accumulates and suppresses DNRR, while the loss of ProT3 in the mutant reduces this uptake, alleviating the inhibitory effect of proline.

**Fig. 2.**
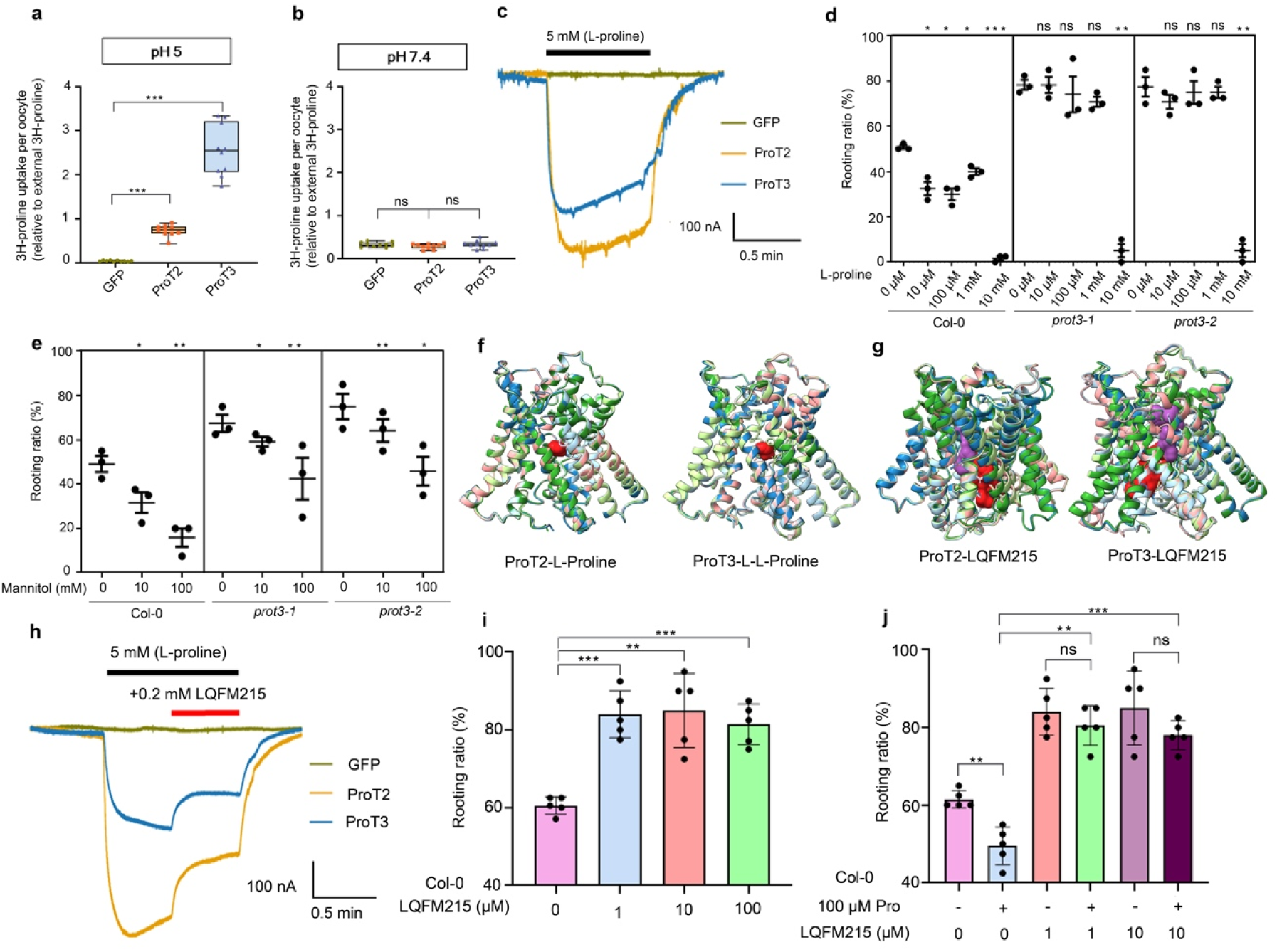
ProT3 modulates DNRR through its proline transport activity and is inhibited by a proline transporter inhibitor LQFM215. a, b, Proline uptake assay in *Xenopus laevis* oocytes. ^3^H-proline uptake was measured at pH 5.0 (a) and pH 7.4 (b). Oocytes expressing ProT2, ProT3, or GFP (negative control) were incubated in kulori buffer containing 0.5 µM ^3^H-proline for 30 min (n = 10 oocytes). c, Electrophysiological recordings of ProT2 and ProT3-mediated proline transport in *Xenopus* oocytes. Representative traces of proline-induced currents recorded from oocytes expressing ProT2, ProT3, or GFP (negative control) at a holding membrane potential of -60 mV at pH5.0. d, Rooting ratio in Col-0 and *prot3* leaf explants treated with L-proline (0-10 mM) (n = 3 biological replicates, 40 explants each). e, Rooting ratio in Col-0 and *prot3* leaf explants under mannitol treatment (0–100 mM) (n = 3 biological replicates, 40 explants each). f, g, Superimposed AlphaFold v3.0.1 models of Arabidopsis ProT2 and ProT3 in complex with L-proline (f) and LQFM215 (g). For ProT2 and ProT3, five predicted models with different colors are superimposed. Model 1, light blue (#a6cee3); model 2, dark blue (#1f78b4); model 3, light green (#b2df8a); model 4, dark green (#33a02c); and model 5, light red (#fb9a99). L-proline is shown in dark red (#e31a1c) in all models. For LQFM215, ligand colors reflect their binding positions. In the ProT2-LQFM215 complex, ligands in models 2 and 4 occupy a similar position and are colored purple (#984ea3), whereas ligands in models 1, 3, and 5 are colored dark red (#e31a1c); the same coloring scheme is applied to the ProT3-LQFM215 complex. h, Electrophysiological analysis showing inhibition of ProT2 and ProT3 by LQFM215. The effect of LQFM215 on ProT2- and ProT3-mediated proline-induced currents was assessed by applying 0.2 mM LQFM215 (red bar) in the presence of 5 mM proline (black bar) at a holding potential of -60 mV at pH5.0. i, Rooting ratio in Col-0 by different concentration of LQFM215 treatment (n = 5 biological replicates, 40 explants each). j, LQFM215 alleviates proline-mediated suppression of DNRR in Col-0 (n = 5 biological replicates, 40 explants each). Data are presented as mean ± s.d.; *n* denotes the number of biological replicates. Statistical significance was determined by two-tailed Student’s *t*-test; *P < 0.05, **P <0.01, ***P < 0.001; ns, not significant.

We next asked whether chemical inhibition of proline transport could mimic the genetic loss of ProT3. Previous studies have shown that LQFM215, a small-molecule inhibitor of mammalian proline transporters (SLC6A7), blocks substrate uptake in synaptosomes ^46,47^. Inspired by this, we hypothesized that LQFM215 might also bind to Arabidopsis ProT2 and ProT3 and inhibit their transport activity. To test this possibility, we first predicted Arabidopsis ProT2/3 interactions with L-proline using AlphaFold v3.0.1. Superimposition of five independent models revealed high-confidence complexes (pTM= 0.87-0.88, ipTM=0.95-0.96 for ProT2; pTM = 0.86, ipTM=0.92-0.94 for ProT3) (Supplementary Table 1), with L-Proline consistently positioned within a conserved central substrate-binding cavity (Fig. 2f). We next generated ligand–protein complexes for LQFM215. Across five independent AlphaFold predictions, LQFM215 adopted multiple partially overlapping orientations in both ProT2 (pTM = 0.84-0.86, ipTM=0.78-0.84) and ProT3 (pTM = 0.82-0.85, ipTM=0.69-0.8), forming a cluster of plausible poses within the substrate pocket (Fig. 2g, Supplementary Table 1), suggesting flexibility in inhibitor engagement with the transporters. Strikingly, nearly all residues predicted to contact L-proline in ProT2 and ProT3 were encompassed within the broader set of residues contacting LQFM215 (Supplementary Table 1). This overlap indicates that LQFM215 occupies—and extends beyond—the endogenous substrate-binding site. Because the inhibitor engages the same core residues required for proline coordination, these structural models strongly suggest a competitive inhibition mechanism in which LQFM215 interferes with the substrate-binding cavity and prevents proline from entering the transport pathway.

Following this *in silico* prediction, we performed electrophysiological recordings in Xenopus oocytes to test whether LQFM215 affects proline transport activity. Application of 0.2 mM LQFM215 markedly reduced the proline-induced inward currents in oocytes expressing either ProT2 or ProT3, indicating an inhibitory effect on transporter activity (Fig. 2h). These results confirmed that LQFM215 functions as an inhibitor of Arabidopsis ProT activity. We further examined the physiological impact of LQFM215 on DNRR in Arabidopsis to evaluate its functional effect in planta. When LQFM215 was applied on leaf explants, it enhanced DNRR by approximately 20% at a concentration as low as 1µM, similar as the enhanced DNRR observed in *prot3* mutants (Fig. 2i). Notably, LQFM215 also rescued the L-proline–induced suppression of DNRR, thereby confirming that its promotive effect on DNRR was mediated through inhibition of proline uptake (Fig. 2j). Taken together, these results demonstrate that compromising proline import, either by knocking out endogenous ProTs function or application of inhibitors of proline transporters, promotes the regeneration of adventitious root from leaf explants.

### *prot3* mutation rescues SA-mediated suppression of DNRR

We next investigated the relationship between SA and proline transporters in regulating DNRR. Exogenous NaSA, an SA mimic, inhibited DNRR in Col-0 ^9^, while *prot3* and *prot2* mutants were insensitive to this suppression (Fig. 3a,b). Similar to what we observed in *prot3* mutants, LQFM215 treatment also rescued the suppression of DNRR caused by exogenously applied SA (Fig. 3c). To test if LQFM215 rescues DNRR defects caused by endogenous SA, we used transgenic plants overexpressing the bifunctional SA synthase gene *Irp9* from the pathogenic bacterium *Yersinia enterocolitica* under the control of the 35S promoter (*35S::Irp9*) ^48^. *35S::Irp9* contains elevated level of SA and SA-derived conjugates and enhances SA-mediated disease resistance ^48^. DNRR was strongly suppressed in *35S::Irp9* (Fig. 3e). However, application of 100 µM of LQFM215 partially restored adventitious root formation from explants of *35S::Irp9* (Fig. 3e), suggesting that interfering proline uptake rescues SA-induced suppression of regeneration. SA also contributes to an age-dependent decline of regeneration ^8, 9^ with reduced DNRR capacity in explants from 8-, 11-, and 14-day old plants. Notably, DNRR in *prot3* mutants remained age-dependent, with the rooting ratio decreasing from 76.2% in 8-day-old explants to 46.4% in 14-day-old explants, despite the overall higher rooting ratio compared to Col-0 (Fig. 3f), suggesting that ProT3 only acts in a branch of SA-mediated suppression of DNRR or other age-dependent signals inhibit DNRR in old leaves. On the other side, L-proline-mediated inhibition of DNRR was also compromised in *NahG* transgenic line which possess reduced SA level due to the expression of a salicylate hydroxylase (Fig. 3g). These data suggested that SA and proline may act synergically to suppress DNRR.

**Fig. 3.**
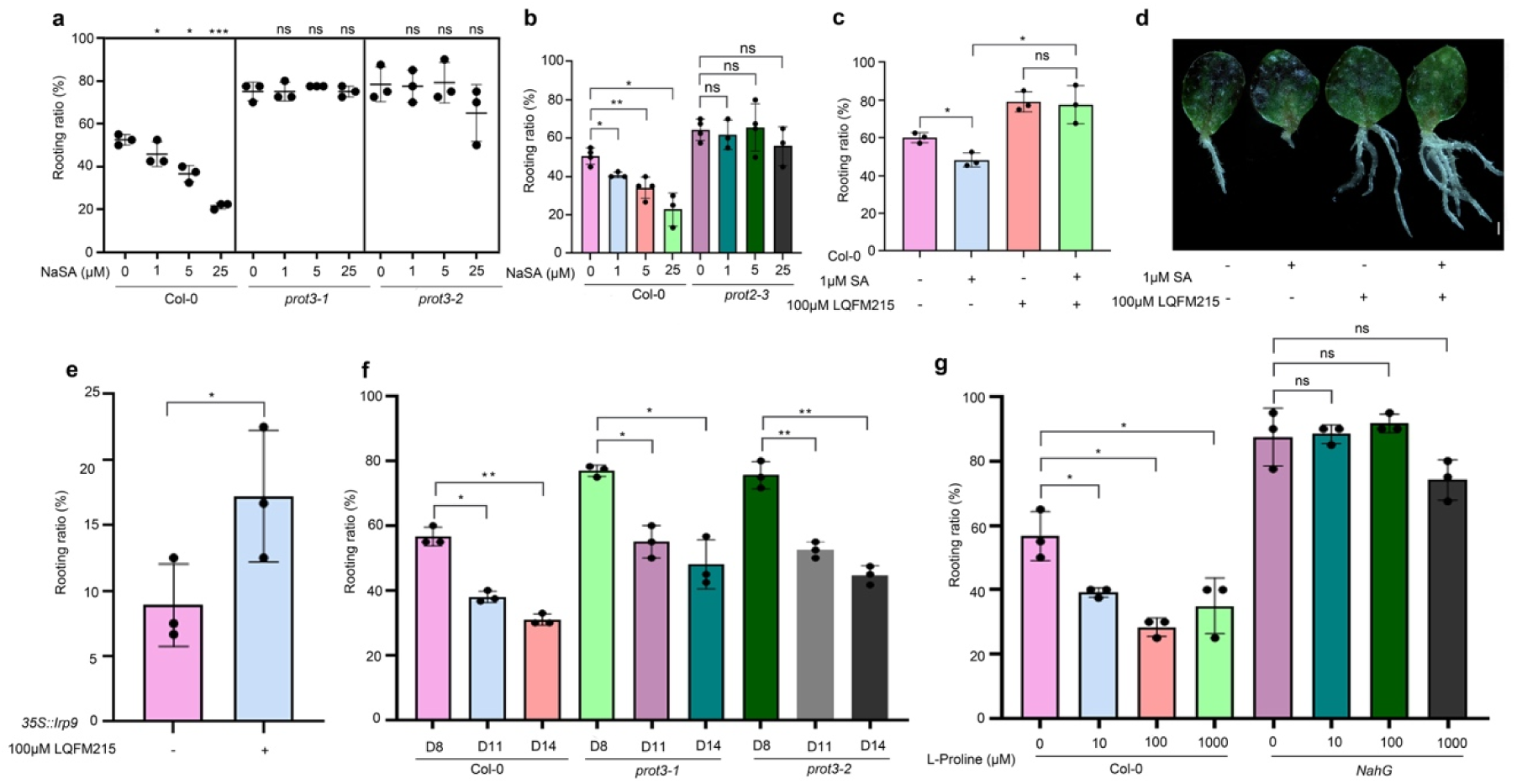
Mutation of ProT3 abolishes salicylic acid (SA)-mediated suppression of DNRR. a, b, Rooting ratio in Col-0, *prot3-1*, *prot3-*2 (a) and *prot2-3* (b) explants treated with NaSA (0–25 µM) (n = 3 biological replicates, 40 explants each). c, Rooting ratio in Col-0 explants co-treated with 1 µM NaSA and 100 µM LQFM215 (n = 3 biological replicates, 40 explants each). d, Representative images showing that 10 µM LQFM215 alleviates SA-mediated suppression of DNRR in Col-0 leaf explants. Scale bar: 1 mm. e, Rooting ratio in *35S::Irp9* (high SA level) with or without 100 µM LQFM215 (n = 3 biological replicates, 40 explants each). f, Rooting ratio in Col-0 and *prot3-2* explants from 8-, 11- and 14-day-old plants (n = 3 biological replicates, 40 explants each). g, Rooting ratio in Col-0 and *NahG* explants treated with L-proline (10 µM–1 mM) (n = 3 biological replicates, 40 explants each). Data presented as mean ± s.d.; n denotes the number of biological replicates. Statistical significance was determined by two-tailed Student’s t-test; *P < 0.05, **P <0.01, ***P < 0.001; ns, not significant.

We further tested whether microbe-induced inhibition of DNRR was affected by ProT3. Treatment with *Pseudomonas syringae pv. tomato* (*Pto*) hrcC- mutants, a compromised bacterium defective in effector delivery but sufficient to induce PTI, inhibited DNRR in *prot3-1* and *prot3-2* mutants to a similar extent as in Col-0 (Extended Data Fig. 2a), indicating that ProT3 does not influence PTI-mediated DNRR suppression. Similarly, 1 µM of flg22, a peptide-derived from bacterial flagellin capable of activating basal immunity, suppressed DNRR in *prot3* as in Col-0 (Extended Data Fig. 2b). These results are consistent with our previous observations that flg22-induced suppression of DNRR was independent of SA ^6^. Taken together, immune response activates multiple pathways to inhibit regeneration, mutation in *prot3* only blocks the SA-mediated suppression of DNRR.

### ProT3 is dispensable for SA-mediated activation of defense genes and inhibition of bacteria growth

SA regulates a tradeoff between regeneration capacity and immunity ^8, 9, 10^. To test whether ProT3 is also required for SA-mediated activation of immunity, we measured the transcript levels of the SA-responsive genes *PR1* and *SARD1* following the same SA treatment used in the SA-mediated suppression of DNRR assay (Fig. 4a,b). The expression level of these SA responsive genes was comparable in Col-0 and *prot3* mutants (Fig. 4a,b), indicating that ProT3 is not required for the activation of these SA responsive genes in 11-day-old seedlings. Previous work showed that the expression of proline transporters including ProT2 and ProT3 were activated upon flg22 treatment, which depletes apoplastic proline as a defense strategy ^26^. We also observed that *prot2* and *prot3* single mutant showed mild enhanced susceptibility to *Pseudomonas syringae* pv. *tomato* DC3000 (*Pto* DC3000) infection (Fig. 4c, Extended Data Fig. 3a). To specifically test if *prot3* is required for SA-mediated suppression of bacterial growth, we monitored bacterial load of *Pto* DC3000 in Col-0 and *prot3* mutants pre-treated with benzothiadiazole (BTH), an SA analogue (Fig. 4c). As expected, the pretreatment of BTH activated SA-mediated immunity and reduced bacterial growth in Col-0 (Fig. 4c). Such reduction was also observed in *prot2* and *prot3* mutants to a similar level (Fig. 4c), further supporting that ProT2 and 3 were dispensable for SA-dependent suppression of bacteria multiplication.

**Fig. 4.**
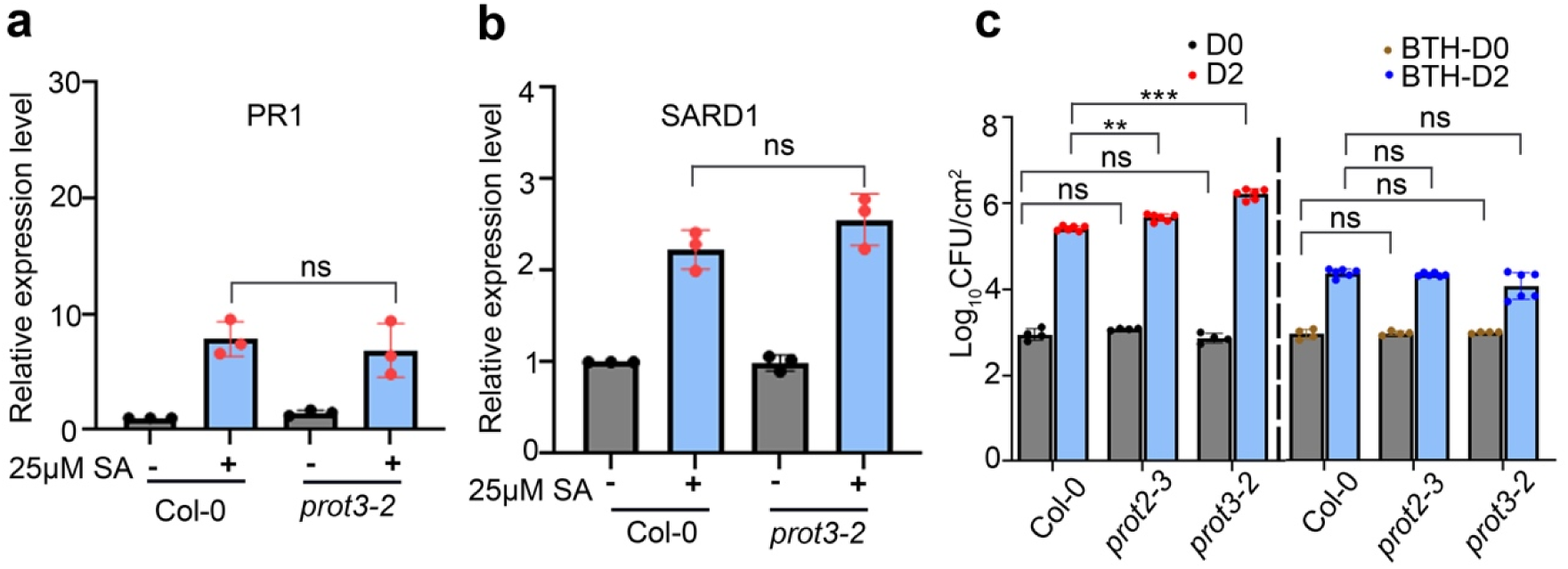
ProT3 does not interfere with SA-induced defense activation. a, b, qRT-PCR analysis of *PR1* (a), *SARD1* (b) expression in Col-0 and *prot3*-2 leaf explants from 11-day old seedlings under 25 μM NaSA treatment. (n = 3 biological replicates). c, Bacterial growth of *Pseudomonas syringae* pv. *tomato* DC3000 in seven-week-old plants of Col-0, *prot2-3*, and *prot3-2*. The left panel shows Pto DC3000 growth kinetics without chemical pretreatment. The right panel shows assessing SA-dependent resistance, where plants were spray treated with 50 µM BTH and inoculated with Pto DC3000 1 day after BTH treatment. Bacterial load was quantified 2 days post inoculation (D2). (n = 4 for D0; n = 6 for D2). Data are presented as mean ± s.d.; Statistical significance was determined by two-tailed Student’s *t*-test; *P < 0.05, **P <0.01, ***P < 0.001; ns, not significant.

### ProT3 interacts with CPK1 to suppress regeneration

When exploring the potential function of ProT3 in regulating regeneration, we identified calcium-dependent protein kinase 1 (CPK1) as a protein interactor of ProT3 from public database (https://thebiogrid.org/) ^49^. Bimolecular fluorescence complementation (BiFC) and co-immunoprecipitation assays based on transiently expressed proteins in *Nicotiana benthamiana* confirmed a physical interaction between ProT3 and CPK1 (Fig. 5a,b). Although *CPK1* expression was not induced by cutting (Extended Data Fig. 3b), two alleles of *cpk1* single mutants showed enhanced DNRR compared to Col-0 (Fig. 5c,d), supporting CPK1 as a negative regulator of regeneration. Similar to *prot3* mutants, *cpk1* mutants also displayed higher WIC formation than Col-0 (Fig. 5e,f), indicating that a ProT3-CPK1 complex cooperatively suppress wound-induced regeneration. CPK1 was also dispensable for SA-mediated protection against *Pto* DC3000 (Fig. 5g) or SA-induced defense gene activation (Fig. 5h,i). However, the *cpk1* mutant did not block SA-mediated suppression of DNRR (Extended Data Fig. 3c), indicating a signaling divergence between ProT3 and CPK1 or redundancy between CPK1 and closely related kinases. Taken together, these molecular and genetic evidence support a conclusion that a ProT3-CPK1 complex is required for wound-induced regeneration but not the activation of SA-mediated defense, decoupling the two key functions downstream of wound-activated SA.

**Fig. 5.**
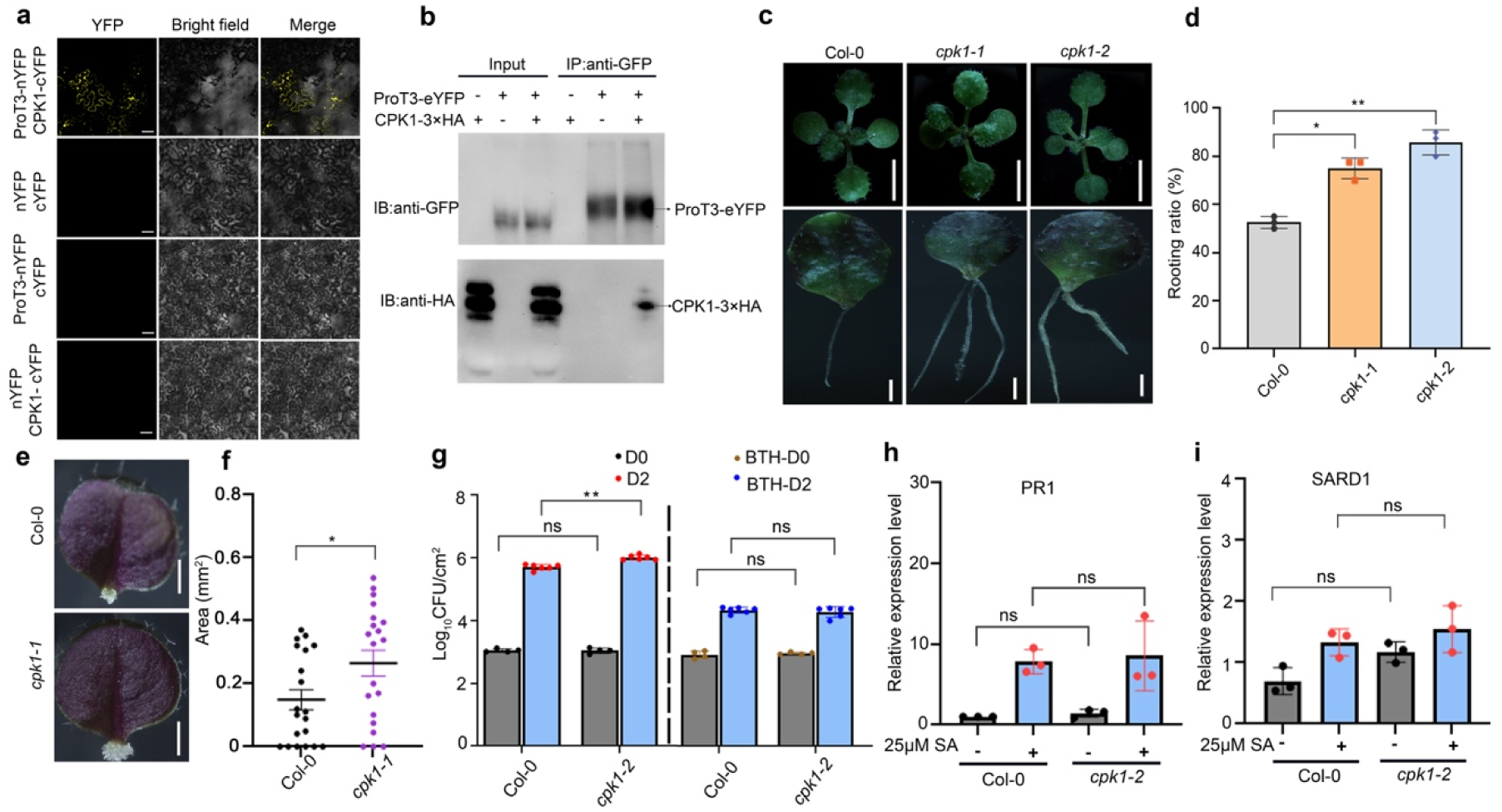
ProT3 interacts with CPK1 to suppress regeneration. a, Bimolecular fluorescence complementation (BiFC) assay showing interaction between ProT3 and CPK1 in *Nicotiana benthamiana*, scale bar, 50 µm. Fluorescent signals were detected 2 days after inoculation with Agrobacteria carrying corresponding plasmids. b, Co-immunoprecipitation assay confirming ProT3-CPK1 interaction in *N. benthamiana*. c, Morphology (top) and DNRR phenotype (bottom) of Col-0, *cpk1-1* and *cpk1-2*, scale bars, 0.5 cm (top) and 1 mm (bottom). d, Rooting ratio in Col-0, *cpk1-1* and *cpk1-2* (n = 3 biological replicates, 40 explants each). e, f, Wound-induced callus (WIC) formation in Col-0, *prot3-2* and *cpk1-1* (e) and quantification of callus area (f), n = 20 petioles per genotype, scale bar, 1 mm. g, Bacterial growth of *Pseudomonas syringae* pv. tomato DC3000 in Col-0 and *cpk1* mutant with or without BTH pretreatment. The left panel shows DC3000 growth kinetics without chemical pretreatment. The right panel shows BTH-induced resistance; plants were spray-treated with 50 µM BTH one day before DC3000 inoculation. Bacterial loads were quantified 2 days after inoculation (n = 4 for D0; n= 6 for D2). h,i, qRT-PCR analysis of *PR1* (h) and *SARD1* (i) expression in 11-day-old Col-0 and *cpk1-2* seedlings treated with 25 μΜ NaSA treatment. (n = 3 biological replicates). Data are presented as mean ± s.d.; Statistical significance was determined by two-tailed Student’s *t*-test; *P < 0.05, **P <0.01, ***P < 0.001; ns, not significant.

### ProT3 regulates ROS equilibrium during DNRR

To interrogate the transcriptomic changes induced by *prot3* mutation, we performed an RNA-seq experiment comparing *prot3* and Col-0 transcriptomes at time 0 (D0) and 1 day after cutting (D1). At D0, we identified 255 and 113 genes that are differentially expressed (DEGs) in blades and petioles, respectively (Fig. 6a). At D1, there were 321 and 194 DEGs between the two genotypes in blades and petioles, respectively (Fig. 6a). It is noteworthy that only 19 DEGs in blades and 8 DEGs in petioles are shared between D0 and D1, indicating a dramatic shift of expression in response to wound (Fig. 6a). We noticed that there was no enrichment of SA-related GO terms in DEGs D0 or D1 by comparing blade or petiole samples from Col-0 and *prot3*, which further supported our observation that *prot3* did not generally contribute to SA-mediated defense (Supplementary Table 2). In addition, key genes involved in DNRR regeneration, including *YUCCA2* and *PLT7*, known to promote regeneration through auxin biosynthesis and meristem maintenance, were upregulated in *prot3* petioles at D1 (Fig. 6b,c), which was consistent with the enhanced adventitious root formation (Fig. 1c,d). To further explore the ProT3-controlled biological processes that are important for regeneration, we performed a hierarchical clustering analysis of all 714 genes that were differentially expressed between *prot3* and Col-0 at either D0 or D1 (Fig. 6b). Notably, in cluster 3, DEGs were more activated in the D1 leaf blades of *prot3* (Extended Data Fig. 4a). In addition, DEGs in cluster 4 were less activated in both blades and petioles of *prot3* (Extended Data Fig. 4b). We noticed that enriched GO terms related to oxygen metabolism such as “response to oxygen-containing compound” in cluster 3 and cluster 4 genes (Supplementary Table 3), consistent with proline’s role as a ROS regulator through its metabolic cycling in cytoplasm and mitochondria. We further assessed ROS accumulation by H2DCFDA staining at cutting sites in Col-0 and *prot3 cpk1* (Fig.6d-g).

**Fig. 6.**
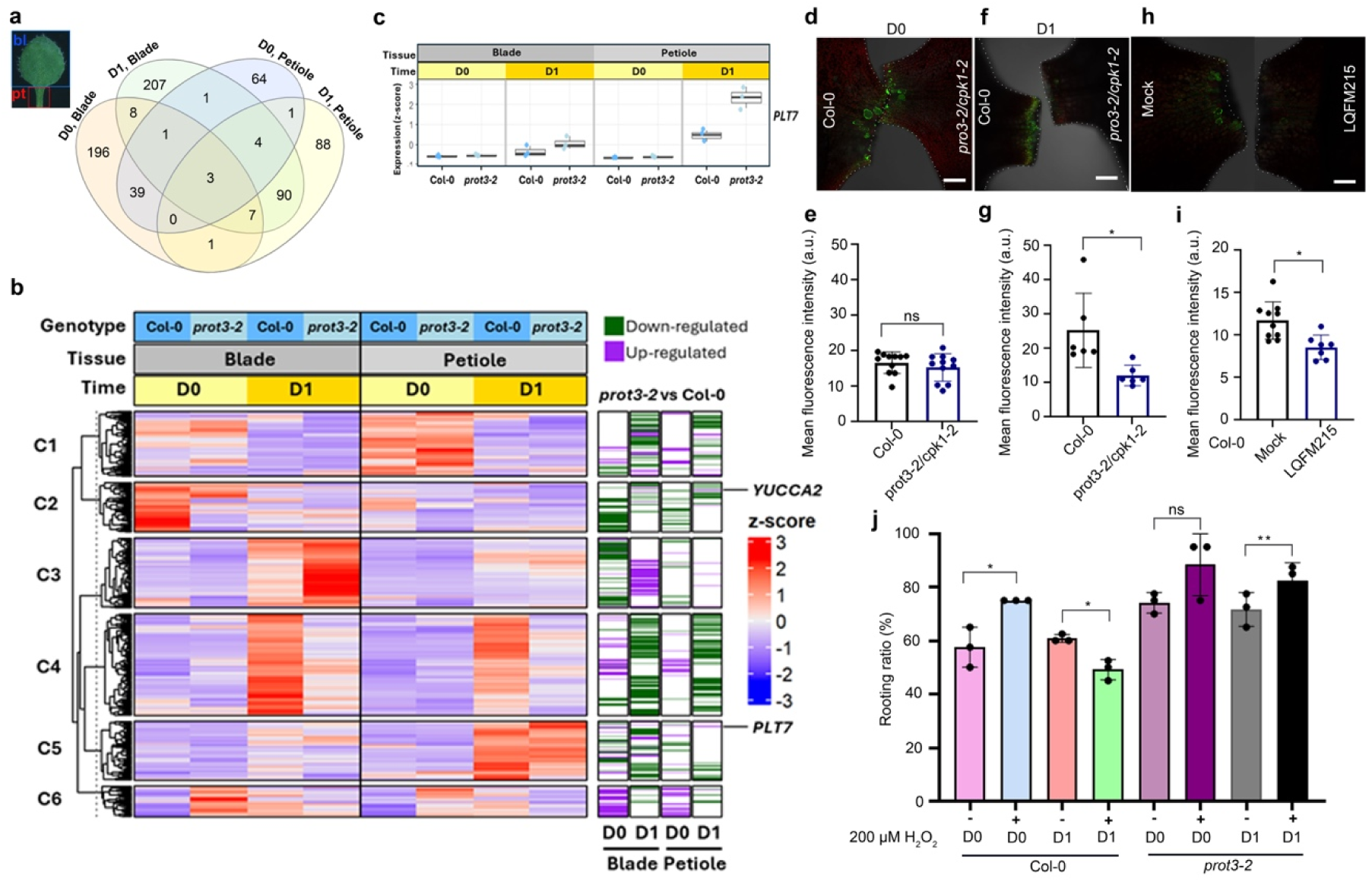
ProT3 regulates reactive oxygen species (ROS) homeostasis during DNRR. a. Arabidopsis leaf showing the regions used for RNA-seq (blade, bl, blue box; petiole, pt, red box) and Venn diagram showing the overlap of differentially expressed genes (DEGs) between *prot3-2* and Col-0 in blade and petiole tissues at D0 and 1 day after cutting (D1). b, Hierarchical clustering of 710 genes that were differentially expressed between Col-0 and *prot3-2* at least of time point in leaf blades or petioles. Statistically significant differentially expressed genes (FDR ≤ 0.05) are marked with green (downregulated) or purple (upregulated) stripes. Representative auxin-related genes *YUCCA2* and *PLT7* are highlighted. c, Expression profiles of *PLT7* in leaf blade and petiole tissues at D0 and 1 day after cutting (D1) in Col-0 and *prot3-2*. Box plots show normalized expression values (z-scores) derived from RNA-seq data. d-g, H2DCFDA fluorescence in leaf explants of Col-0 and *prot3-2/cpk1-2* mutants at D0 (d) and 1DAC (f). Quantification of H2DCFDA fluorescence was summarized below representative images. Dotted lines outline wound sites. scale bar: 200 μm. h,i, H2DCFDA fluorescence in leaf explants of Col-0 treated with 10 μM LQFM215 treatment at 1DAC. Quantification of fluorescence intensity is shown in (i). j, Effect of exogenous H₂O₂ treatment on DNRR in Col-0 and *prot3-2* at different times after cutting. H_2_O_2_ was applied either immediately after cutting (D0) or at D1. Rooting ratios from each treatment are shown. (n = 3 biological replicates, each with 40 explants). Data are presented as mean ± s.d.; Statistical significance was determined by two-tailed Student’s *t*-test; *P < 0.05, **P <0.01, and ***P < 0.001; ns, not significant.

ROS signal was immediately accumulated at wound sites after cutting, no difference was observed between Col-0 and *prot3-2cpk1-2* double mutant at D0 (Fig. 6d,e). However, at D1, the double mutants showed reduced ROS accumulation at cutting sites compared to Col-0 (Fig. 6f,g). Application of 10 µM LQFM215 immediately after cutting also reduced ROS accumulation at D1 (Fig. 6h,i). Previous studies have suggested that wound-induced ROS signal promotes DNRR ^50^. Consistently, we also observed that H₂O₂ treatment at time 0 promoted DNRR in Col-0 (Fig. 6j). In contrast, application H_2_O_2_ at 1 day after cutting reduced rooting by ∼15% in Col-0 (Fig. 6j), suggesting that a reduced sensitivity to, or level of, ROS signal at later stage is required for full regeneration capacity. Strikingly, *prot3* mutants were insensitive to both the promotive effect of early H₂O₂ treatment or the inhibitory effect of H_2_O_2_ applied at D1 (Fig. 6j), indicating that loss of the ProT3-CPK1 complex facilitate a permissive ROS environment at 1 day after cutting for root regeneration. While wound-induced ROS accumulation is required to initiate regeneration ^50^, subsequent ROS reduction may be necessary for DNRR progression. Together, these results suggest that the ProT3–CPK1 complex modulates ROS dynamics during DNRR, integrating early wound-induced stress signals into regenerative programs.

### LQFM215 promotes regeneration in crops

The *ProT* gene family is widely present and highly conserved in plants (Extended Data Fig. 5). We further tested the effect of LQFM215 on promoting regeneration in tomato, tobacco and rice. In tomato (*Solanum lycopersicum* cv. Money Maker), treatment of hypocotyl explants with LQFM215 significantly increased callus size on callus-induction medium (CIM) compared with mock treatment (Fig. 7a,b). Similarly, treatment of leaf explants with LQFM215 resulted in a higher number of adventitious roots per explant, with all root classes increased relative to mock (Fig. 7c,d). In *N. benthamiana*, leaf disc explants of uniform size treated with LQFM215 exhibited higher shoot regeneration efficiency and more shoots per explant compared with controls (Fig. 7e,f). In rice (*Nipponbare*), scutellum-derived calli cultured on CIM were larger upon LQFM215 treatment, indicating enhanced callus proliferation (Fig. 7g,h).

**Fig. 7.**
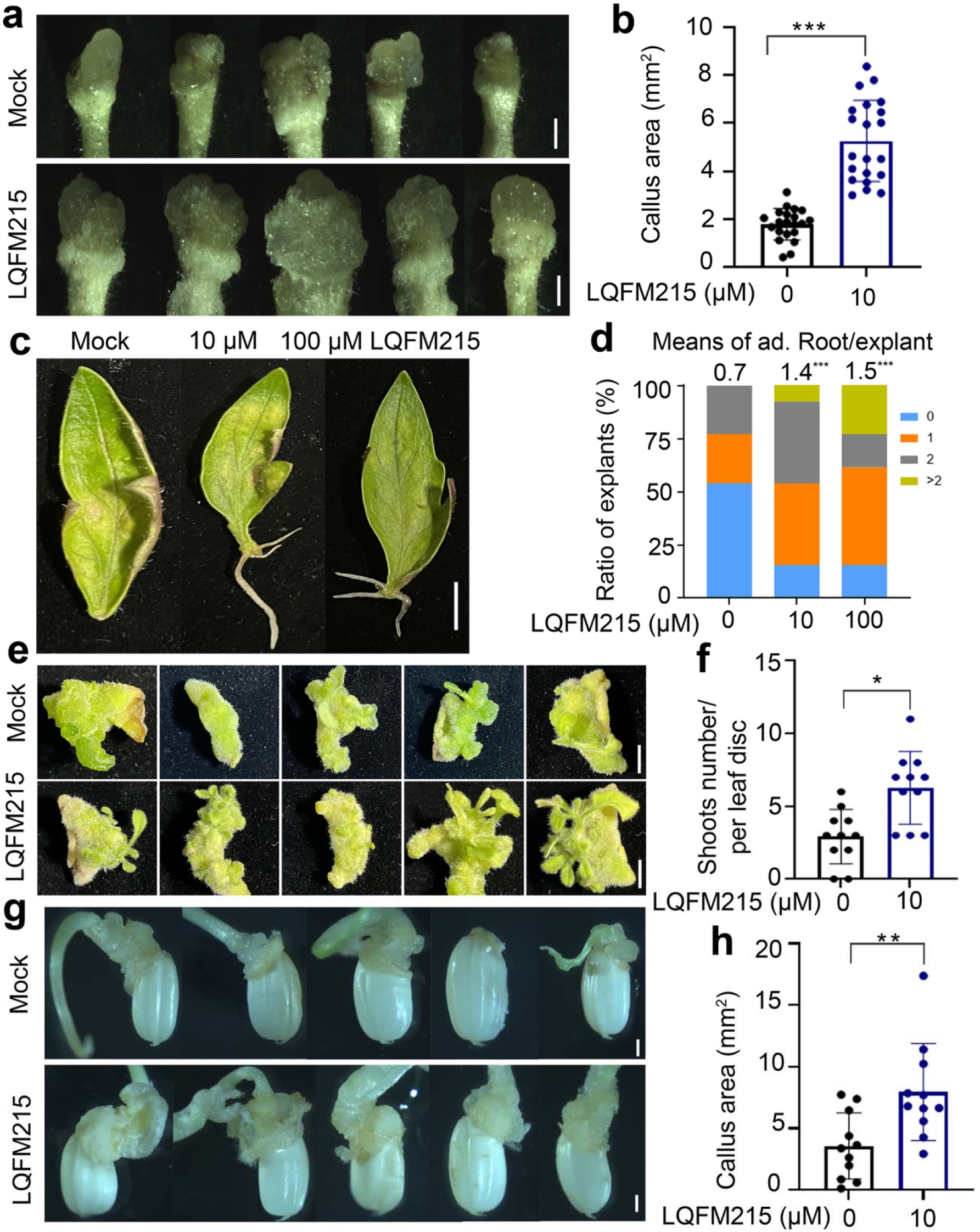
LQFM215 enhances regeneration across multiple crop species. a, b, Tomato leaf explants treated with LQFM215 (0, 10, 100 µM) showing enhanced adventitious root formation at day 10 (a) and distribution of explants according to the number of adventitious roots per explant, expressed as a percentage of total explants (b, n = 20); Mann–Whitney U test; scale bar, 5 mm. c, d, Tomato hypocotyl-derived callus formation with or without 10 µM LQFM215 (c) and quantification (d, n = 21 explants); scale bar, 1 mm. e, f, Shoot regeneration from *Nicotiana benthamiana* leaf discs with or without 10 µM LQFM215 (e); shoot counts after 30 d (f, n=11); scale bar, 5 mm. g, h, Scutellum-derived callus formation from rice (*Oryza sativa* cv. Nipponbare) seeds with or without 10 µM LQFM215 (g) and quantification (h, n = 11 seeds); scale bars, 1 mm. Unless otherwise indicated, data are presented as mean ± s.d.; n denotes the number of biological replicates. Panels showing percentage distributions (Fig. 7d) represent data from a single experiment without error bars. Statistical significance was determined by two-tailed Student’s t-test; *P < 0.05, **P <0.01, and ***P < 0.001; ns, not significant.

Collectively, these results demonstrated that LQFM215 enhances multiple modes of regeneration across diverse crops, extending its function beyond Arabidopsis. This raises the possibility that LQFM215 might help overcome the trade-off between disease resistance and regenerative capacity, providing a strategy to improve regeneration in elite crop varieties while maintaining desirable agronomic traits.

## Discussion

Our study identifies ProTs as critical regulators of DNRR in *Arabidopsis*. We uncovered their role in balancing SA-mediated regeneration and immunity. ProTs are required for SA-mediated suppression of regeneration but not for defense. The function of ProTs is probably achieved via a complex with CPK1. Notably, LQFM215, an inhibitor of plant ProTs, is sufficient to alleviate SA-mediated suppression of DNRR in multiple plant species, suggesting that proline metabolism represents a conserved mechanism linking biotic and abiotic stresses to regenerative capacity. These findings not only broaden the known functions of proline transporters beyond abiotic stress tolerance but also provide new insights into the molecular basis of the growth–defense trade-off in plants.

Wound-induced ROS signaling plays a key role in wound healing and organ regeneration across both animals and plants ^51, 52, 53^. The oxidative burst serves as an early cue that guides wound closure, immune cell recruitment, and stem/progenitor activation. In zebrafish tail regeneration, ROS recruit Hedgehog-expressing cells to the wound site, thereby activating Hedgehog signaling essential for regeneration ^54^. In planarians, ROS acts upstream of the MAPK/ERK cascade and can even rescue regeneration when ERK signaling is inhibited ^55^. In amphibians such as axolotl, H₂O₂ contributes to blastema formation and growth by activating regeneration-related genes and recruiting immune cells to the injury site ^56, 57^. In plants, early ROS signals, likely produced directly by tissue damage, play a promotive role in regeneration ^50, 58, 59, 60, 61, 62^. However, evidence also indicates that excess ROS can inhibit shoot regeneration ^63, 64^ and microcallus formation from protoplasts ^65^. Moreover, genetic reduction of ROS levels enhances DNRR or shoot regeneration in Arabidopsis ^58, 66^. Together, these observations suggest that while ROS act as essential early wound signals, sustained or excessive ROS accumulation may exert inhibitory effects on regeneration beyond their initial signaling role.

Our results showed that the effect of exogenous H₂O₂ switched from promotive to inhibitory within one day after cutting, and that mutation in ProT3 blocked this sensitivity (Fig. 6j), suggesting that ProT3 may play a role in regulating endogenous ROS dynamics or cellular responses to ROS. In mitochondria, proline dehydrogenase (ProDH) catalyzes the oxidation of proline to pyrroline-5-carboxylate (P5C), transferring electrons to the mitochondrial electron transport chain ^67, 68, 69^. This reaction generates ROS, primarily superoxide (O₂•⁻) and hydrogen peroxide (H₂O₂), which serve as signaling molecules that activate downstream defense and stress-response pathways ^70^. Following leaf detachment, AtProDH1 was rapidly induced within one hour in explants, exhibiting a pattern similar to that of ProT3 (Extended Data Fig. 6). These observations suggest that proline transport and degradation may be coordinately regulated to promote wound-induced ROS production. We speculate that excessive or prolonged ROS accumulation derived from proline catabolism—as observed one day post-cutting at wound sites—may inhibit the development of root founder cells, thereby constraining DNRR. Previous work showed that ROS levels increased between 3–6 days after calli were cultured on shoot induction medium (SIM), then declined from days 8 to 16. In mutants of *dcc1*, a thioredoxin, and *ca2* (carbonic anhydrases 2), an interactor of *dcc1*, ROS levels in calli remained high through day 16, correlating with a reduced shoot regeneration phenotype ^66^. In contrast, in the *prot3 cpk1* mutants, a reduced ROS level at later stages post-wounding may create a permissive environment for root founder cell establishment (Fig. 6f,g). The dynamics of ROS levels after wounding and their roles in wound-induced regeneration may explain dosage- and timing dependent phenotype of ROS impact on various forms of plant regeneration. Interestingly, *ProT3* is activated at wound sites but later excluded from developing callus (Fig. 1i), suggesting that a low proline, probably leading to a low ROS condition, is necessary for proceeding with the adventitious root initiation. Thus, proline transporters such as ProT3 and ProT2 may act as metabolic gatekeepers that couple amino acid flux to redox dynamics. Evidence from plant immunity revealed that SA induces *ProDH* expression during the hypersensitive response, and ProDH activity is required for full ROS accumulation and pathogen resistance induced by SA ^70^. It is possible that the loss of ProTs rescued SA-mediated suppression of DNRR by counteracting the SA-induced ROS level. Notably, other amino acid transporters, including LHT1 ^26^ and members of the AAP family ^28^, may also contribute to proline homeostasis in addition to ProTs. Beyond its role as a ROS regulator, proline may also serve as a metabolic substrate or osmo-regulator during regeneration. Although our data show that ProT2/3 transport proline and that their function is inhibited by LQFM215, we cannot rule out the possibility that other ProT cargos also contribute to regeneration. How proline flux and subcellular localization are precisely regulated to maintain redox balance remains a key question for elucidating the mechanistic link stress response and regenerative capacity.

In plants, regeneration and immunity are tightly interconnected processes. Elite crop varieties bred for durable disease resistance often exhibit constitutive or primed immune activity, including elevated salicylic acid (SA) signaling, stronger microbe-associated molecular pattern (MAMP) responses, and enhanced hypersensitive cell death. While these traits confer resilience against pathogens, they may also compromise regenerative competence ^6^. For example, IR64, a high-yielding indica rice variety carrying resistance genes such as Xa4 for bacterial blight, has been widely used in breeding programs but remains one of the most recalcitrant cultivars for regeneration and Agrobacterium-mediated transformation, requiring specialized tissue culture protocols for plant recovery ^71, 72^. Although there is no direct evidence that recalcitrance is caused by enhanced defense responses, strategies that transiently suppress defense signaling or fine-tune immune activation during plant tissue culture or de novo organogenesis could help overcome these limitations. Notably, we showed that pharmacological inhibition of proline transport by LQFM215 alleviated SA-induced suppression of regeneration in multiple plant species, indicating that proline flux functions as a tunable regulatory node in the antagonistic relationship between plant immunity and regeneration.

## Methods

### Plant growth and transformation

*Arabidopsis thaliana* T-DNA lines and transgenic lines used in this study are in the Columbia-0 (Col-0) ecotype. Homozygous lines were selected by gene specific primers LP and RP, in combination with T-DNA vector primers (Supplementary Table 4). Primers used for vector construction are listed in Supplementary Table 4. Double knockout lines were generated by crossing homozygous single *prot3-2* with *prot2-3* to form the *prot2-3prot3-2* double mutant, *prot3-2* with *cpk1-2* to form the *prot3-2cpk1-2* double mutant. Transgenic constructs were introduced via Agrobacterium tumefaciens-mediated transformation using the floral dip method ^73^. Additionally, *Nicotiana benthamiana* plants were cultivated under long-day conditions (16 h of light / 8 h of dark) in the growth room at University of Georgia, maintained at 22 ℃, and 70% relative humidity, with a light intensity of 30000 lumens/m^2^.

### Characterization of the mutant DNRR phenotype and wounding-induced callus formation

DNRR phenotyping was performed as previously described ^74^, with minor modifications. Arabidopsis seeds for DNRR testing were surface sterilized by 70% ethanol, germinated, and grown on ½ MS media (Murashige & Skoog Basal Medium with Vitamins, PhytoTechnology Laboratories) which were placed in the growth chambers (Percival, USA) with continuous light at 22°C. The 1st and 2nd true leaves from 11-day-old seedlings were cut at the junction between the leaf blade and petiole and placed with the abaxial side down onto Gamborg’s B5 media (RPI Research Products International). Explants were kept in continuous light condition. The adventitious roots were counted every day under a dissecting microscope starting from 6 DAC until the 14 DAC day as previously described ^9^. Morphological assessments of whole plants were documented using a Canon EOS Rebel T7 DSLR Camera (EF-S 18–55mm f/3.5–5.6 IS II). Leaf explants with adventitious root formation were photographed using a Zeiss Axio Zoom.V16 Stereo Microscope. DNRR assays performed on Phytagel plates and sand plates (for *Pto* DC3000 hrC^-^ treatments) were performed as previously outlined ^11^. The callus size was scored 10 days after excision and callus areas were quantified using ImageJ software.

### Chemical treatments

To investigate the effects of various chemical cues on DNRR, detached Arabidopsis leaf explants were subjected to the following treatments.

For salicylic acid (SA) treatments, sodium salicylate (NaSA) was added to B5 medium at final concentrations of 0 μM (mock), 1 μM, 5 μM, and 25 μM. Explants were transferred onto the respective media, and the adventitious root formation ratio was quantified as described above.

For the regeneration response to proline, detached leaf explants were transferred onto B5 medium containing different concentrations of L-proline (0 μM, 10 μM, 100 μM, 1 mM, and 10 mM).

For testing the DNRR response to flg22 elicitor treatment, detached leaf explants were pre-treated by spraying them with 1 μM flg22 for 1 hour. The explants were then placed on B5 medium, and 10 µl of flg22 was applied directly to the wound sites after cutting.

For osmotic stress assays, D-mannitol was added to B5 medium at final concentrations of 0 mM (mock), 10 mM, and 100 mM to impose mild and moderate osmotic stress conditions, respectively.

For ROS treatments, freshly prepared 200 μM H₂O₂ was directly applied to the wound sites of leaf explants at different time points after cutting to evaluate regeneration responses under oxidative stress.

To test whether LQFM215, a small-molecule inhibitor of proline transporters, promotes regeneration, LQFM215 was applied topically to the wound sites of leaf explants immediately after placement on the regeneration medium. Unless otherwise indicated, 3–10 μL of 1, 10 or 100 μM LQFM215 was pipetted directly onto each cutting site. The explants were subsequently cultured under standard regeneration conditions, and regeneration phenotypes were evaluated at the indicated time points.

### qRT-PCR analysis

For quantitative reverse transcription PCR (qRT-PCR) analysis, total RNA was extracted from leaf blade cuttings at different time points using the E.Z.N.A Plant RNA Kit (R6827-01, Omega BIO-TEK). First-strand cDNA was synthesized from total RNA using GoScript™ Reverse Transcriptase (A5003, Promega). Real-time RT-PCR was conducted with iQ SYBR Green Supermix (Bio-rad) on the Applied Biosystems QuantStudio 5 real-time PCR system (Applied Biosystems by Thermo Fisher Scientific). qRT-PCR procedures and data analysis were performed as previously described ^9^. Primers used for qRT-PCR are listed in Supplemental Table 4. The endogenous control *Tublin 2* was not differentially expressed from time 0 to 2 days after cutting in explants ^45^. It is also not affected by the genotypes ^45^.

### GUS reporter assay for *ProT* genes expression

Plasmids *pKGWFS7-ProT1Pro::GUS*, *pKGWFS7-ProT2Pro::GUS*, *pKGWFS7-ProT3Pro::GUS* and *pKGWFS7-CPK1Pro::GUS* were constructed to examine tissue-specific expression of ProTs and CPK1, with the GUS marker protein driven by the 2576 bp, 2465 bp, 1919 bp and 2123 bp promoter regions of *ProT1*, *ProT2*, *ProT3*, and *CPK1* respectively. GUS activity was visualized by staining young seedlings and leaf blade cuttings at different time points from various transgenic lines overnight in 5-Bromo-4-chloro-3-indolyl-b-D-glucuronide (X-Gluc), as previously described ^9^. Subsequently, tissues were cleared in 70% (v/v) ethanol. Cleared samples were observed and photographed using a Zeiss Axio Zoom.V16 Stereo Microscope and processed using Zeiss ZEN 3.8 and ImageJ software.

### Bimolecular fluorescence complementation (BiFC) of ProT3 and CPK1

For BiFC analysis, the full-length coding sequences (CDS) of *ProT3* and *CPK1* were amplified and were subcloned into the pENTR™/D-TOPO™ Cloning Kit (Thermal fisher scientific) to form the entry vetor *pENTR-ProT3* and *pENTR-CPK1* respectively. The resultant plasmids were then introduced to the tagging vectors *pUBN-nYFP* and *pUBN-cYFP* by using Gateway™ LR Clonase™ II Enzyme mix Kit (Thermal fisher scientific) to generate *ProT3-nYFP* and *CPK1-cYFP* constructs. The primers used for vector construction are listed in Supplementary Table 4. The resulting plasmids were introduced into *Agrobacterium tumefaciens* strain GV3101 by chemical transformation. Single colonies were cultured overnight in LB medium containing appropriate antibiotics, harvested by centrifugation, and resuspended in infiltration buffer (10 mM MgCl₂, 10 mM MES, pH 5.6, and 150 µM acetosyringone) to an OD₆₀₀ = 0.4. The suspensions carrying *ProT3-nYFP* and *CPK1-cYFP* were mixed at a 1:1 ratio before infiltration. The Agrobacterium mixtures were infiltrated into the abaxial side of 4–5-week-old *Nicotiana benthamiana* leaves using a needleless syringe. After infiltration, plants were maintained in the dark for approximately 48 h to facilitate transient expression. YFP fluorescence signals were first detectable around 40–48 h post infiltration, and the strongest fluorescence was typically observed at 48–60 h. Fluorescence was examined using a Zeiss LSM880 confocal laser scanning microscope with excitation at 514 nm and emission collected between 525–560 nm.

### Co-immunoprecipitation of ProT3 and CPK1

For co-immunoprecipitation (Co-IP) assays, entry vetor *pENTR-ProT3* and *pENTR-CPK1* were cloned into the binary vectors pGWB641 (driven by the CaMV 35S promoter, C-terminal EYFP tag) and pGWB614 (35S promoter, C-terminal 3×HA tag), respectively. The resulting constructs (*35S:ProT3-EYFP* and *35S:CPK1-3×HA*) were introduced into *Agrobacterium tumefaciens* strain GV3101 by chemical transformation. Single colonies were cultured overnight in LB medium with appropriate antibiotics, harvested by centrifugation, and resuspended in infiltration buffer (10 mM MgCl₂, 10 mM MES, 150 µM acetosyringone, pH 5.6) to an OD₆₀₀ of 0.1. Equal volumes of Agrobacterium suspensions carrying ProT3-EYFP and CPK1-3×HA were mixed (1:1 ratio) and infiltrated into 4–5-week-old *Nicotiana benthamiana* leaves using a needleless syringe. Infiltrated plants were maintained in darkness for 2 days to allow transient protein expression. To prevent protein degradation, 50 µM MG-132 (Sigma-Aldrich) was infiltrated into the same leaf areas 24 h post Agrobacterium infiltration, and leaves were harvested 48 h post infiltration for protein extraction.

Total protein was extracted from infiltrated *Nicotiana benthamiana* leaves with protein extraction buffer (50 mM Tris-HCl (pH 7.5), 100 mM NaCl, 1 mM EDTA, 10 mM NaF, 5 mM Na3VO4, 0.25% Triton X-100, 0.25% NP-40, 1 mM PMSF, 1× protease inhibitor cocktail), and then incubated with 25 μl anti-GFP agarose beads (Chromotek, gta-20) for 3 h at 4 °C. The beads were washed 3 times with protein extraction buffer, and the precipitated proteins were eluted with 2× SDS loading buffer at 95 °C for 5 min. The samples were subjected to immunoblot analysis using anti-GFP and anti-HA through western blot respectively.

### RNA-seq analysis

Young leaf blades and petioles from Col-0 and *prot3* plants were collected at D0 and D1 (1 day after cutting), with approximately 20 leaf blades per genotype per time point for each biological replicate. Three biological replicates were prepared for each genotype/treatment. Samples were frozen in liquid nitrogen and milled using a homogenizer (OMNI International). Total RNA was extracted using the E.Z.N.A Plant RNA Kit (R6827-01, Omega BIO-TEK), and RNA concentration was measured using a Nanodrop spectrophotometer (Thermo Scientific). RNA samples were sent to Admera Health (https://www.admerahealth.com) for library preparation and sequencing. RNA integrity was assessed using TapeStation or BioAnalyzer High Sensitivity RNA Assays (Agilent Technologies), and RNA was quantified using an Infinite F Nano+ 200 Pro (Tecan, Switzerland). Poly(A)+ RNA was purified and used for strand-specific library construction with the NEBNext® Ultra™ II Directional RNA Library Prep Kit for Illumina®, following the manufacturer’s instructions. Equimolar pooling of libraries was performed based on QC values and sequenced on an Illumina NovaSeq X Plus platform with a read length configuration of 150 PE for 40 M PE reads per sample (20M in each direction).

Sequencing data processing, including read mapping, alignment, and analysis of differentially expressed genes (DEGs) were performed as previously described ^75^. Differential gene expression analysis was conducted in R using the edgeR package ^76^ to compare the two genotypes (Col-0 vs *prot3*) across different tissues (leaf blade or petiole) at each time point (D0 or D1). Genes with very low expression levels were excluded by retaining only those with at least 1 Transcripts Per Million (TPM) in five or more samples. The Benjamini–Hochberg correction was applied to adjust p-values for multiple testing ^77^. Genes were defined as differentially expressed when the false discovery rate (FDR) was ≤ 0.05 and the fold-change was at least twofold. Gene Ontology (GO) enrichment analyses were performed at the Plant GeneSet Enrichment Analysis Toolkit (PlantGSEA) online webserver ^78^. Hierarchical clustering of expression data was conducted using the ComplexHeatmap package in R^79^, applying the Euclidean distance and the complete linkage method. Expression levels, expressed as TPM values, were standardized to z-scores prior to clustering. Raw RNA-seq reads are deposited in the NCBI Sequence Read Archive (SRA) with BioProject ID PRJNA1357857.

### Proline transport assay in *Xenopus laevis* oocytes

The coding DNA sequences of *ProT2* and *ProT3* were cloned into *Xenopus* expression vector pNB1u using USER cloning ^80, 81^ Linear DNA templates for *in vitro* transcription were generated by PCR from the pNB1 plasmid using forward primer (5’–AATTAACCCTCACTAAAGGGTTGTAATACGACTCACTATAGGG–3’) and reverse primer (5’–TTTTTTTTTTTTTTTTTTTTTTTTTTTTTATACTCAAGCTAGCCTCGAG–3’) and a Phusion High-Fidelity DNA Polymerase (Thermo Fisher Scientific) following the manufacturer’s instructions. The PCR product was purified using E.Z.N.A. Cycle Pure Kit (Omega BIO-TEK) following the manufacturer’s instructions. Capped cRNA was in vitro synthesized using mMessage mMachine T7 Kit (InVitrogen), following the manufacturer’s instructions. The cRNA concentration was normalized to 500 ng µl^-1^ for oocyte injection.

Defolliculated *Xenopus laevis* oocytes at stage V–VI (Ecocyte Bioscience, Germany) were microinjected with 50.6 nL of cRNA using the Drummond NANOJECT II (Drummond Scientific Company, Broomall, Pennsylvania, USA), with GFP serving as a negative control. The injected oocytes were incubated at 16°C for three days in HEPES-based kulori buffer (90 mM NaCl, 1 mM KCl, 1 mM MgCl₂, 1 mM CaCl₂, 10 mM HEPES, pH 7.4) supplemented with amikacin (100 µg mL⁻¹).

Three days after cRNA injection, oocytes were initially pre-incubated for 5 min in kulori buffer pH 5.0 (90 mM NaCl, 1 mM KCl, 1 mM MgCl₂, 1 mM CaCl₂, 10 mM MES, pH 5) or pH 7.4 (90 mM NaCl, 1 mM KCl, 1 mM MgCl₂, 1 mM CaCl₂, 10 mM HEPES, pH 7.4) and then transferred to 150 µl of the same buffer (pH 5.0 or pH 7.4) containing 47 µCi/ml of ³H-L-proline (93.2 Ci/mmol, Revvity) and incubated for 30 min. Afterwards, the oocytes were washed four times with kulori buffer of the respective pH. Each oocyte was then placed in a scintillation vial and lysed in 100 µl of 10% (w/v) SDS by vigorous vortexing. Subsequently, 2.5 ml of EcoScint^TM^ scintillation fluid (National Diagnostics) was added to the vial, mixed thoroughly by vortexing, and the radioactivity was measured using liquid scintillation counting.

### Two-electrode voltage-clamp electrophysiology

The two-electrode voltage-clamp (TEVC) technique was employed to determine electrogenicity of the proline transport via ProT2 and ProT3 and further characterize them. The TEVC recordings were carried out using the automated Roboocyte2 system (Multichannel Systems, Reutlingen, Germany). The electrodes (resistance 280-1000 kΩ) were filled with a solution containing 3 M potassium chlorideand 1.5 M potassium acetate. Oocytes expressing ProT2, ProT3, or GFP (negative control) were clamped at - 60 mV membrane potential and currents were recorded under continuous perfusion with MES-based ekulori buffer (2 mM LaCl3, 90 mM NaCl, 1 mM KCl, 1 mM MgCl2, 1 mM CaCl2, 10 mM MES pH 5), with or without 5 mM L-proline. To test the inhibitory effect of LQFM215, 0.2 mM LQFM215 and 5 mM L-proline were used.

### Bacterial growth assay

Bacterial growth curve assays were performed as previously described ^75^. Briefly, *Pseudomonas syringae pv. tomato* (*Pto*) DC3000 strain was suspended in 10 mM MgCl₂ to a final concentration of 1 x 10LJ CFU/mL and infiltrated into Arabidopsis leaves using a 1 mL syringe without a needle. After inoculation, plants were covered with transparent lids for one hour. Day 0 samples were collected immediately after lid removal, and Day 3 samples were collected three days post-infiltration (dpi). Each sample consisted of four leaf discs (one from each of four individual leaves) punched with a uniform-sized leaf corer. The discs were homogenized using a homogenizer (OMNI International), and the resulting homogenate was serially diluted (10⁻¹ to 10⁻LJ). A 10 μL aliquot of each dilution was plated onto King’s B solid medium containing rifamycin and cycloheximide and incubated at room temperature for two days. Colony-forming units (CFUs) were manually counted and normalized to the total leaf area sampled.

For benzothiadiazole (BTH) pretreatment, an SA analogue, plants were sprayed with 50 μM BTH solution using a sterilized sprayer one day before pathogen inoculation. After spraying, plants were kept in the greenhouse overnight under the same growth conditions. The following day, bacterial growth curve assays were conducted as described above.

### Reactive oxygen species (ROS) detection

Reactive oxygen species were also visualized using the fluorescent probe 2’,7’-dichlorodihydrofluorescein diacetate (H₂DCFDA; D399, Thermo Fisher Scientific) as previously described with minor modification ^82^. A 1 mM H₂DCFDA stock solution was prepared in DMSO and stored at −20 °C in the dark. Before use, the stock was diluted with 10 mM Tris-HCl buffer (pH 7.2) to a working concentration of 10 µM. Leaf explants were collected at indicated time points (D0 and 1 days after cutting) and immersed in 10 µM H₂DCFDA solution, vacuum-infiltrated at 60 kPa for 5 min, and incubated at room temperature for 10 min in the dark. After staining, samples were rinsed thoroughly with 10 mM Tris-HCl buffer to remove residual dye. Fluorescence was observed using a Zeiss LSM 880 confocal laser scanning microscope with excitation at 488 nm and emission at 530 nm. Green fluorescence indicated ROS accumulation, while red autofluorescence corresponded to chlorophyll. Fluorescent areas were quantified using ImageJ software by measuring the average signal intensity in a defined area. The area was defined as tissues 2 mm away from the cutting edge.

### Regeneration assays in crop species with LQFM215

For tomato DNRR, *Solanum lycopersicum* cv. Money Maker seeds were sterilized in 70% ethanol for 30 s, followed by 10% bleach for 15 minutes twice, rinsed five times with sterile water, dried on sterile filter paper, and germinated on ½ MS at 22 °C under continuous light for ∼2 weeks. The first true leaves were excised at the petiole fully using sterile scissors and placed adaxial side up on B5 plates. Apply 3 µL of 10 µM and 100 µM of LQFM215 directly onto the wound sites. Explants were incubated at 22 °C, continuous light, and adventitious roots (AR) were scored at 10 days post-cutting.

For hypocotyl-callus inducion, 10-day-old tomato seedlings were cut with scissors and hypocotyl segments were incubated on CIM (MS + 3% sucrose, 2 mg/L 2,4-D, 0.1 mg/L 6-BA, 0.375% Phytagel; pH 5.8). Immediately after placement, 3 µL of 10 µM LQFM215 was applied to the cut surfaces. Plates were kept at 22 °C, continuous light, for ∼2 weeks before callus phenotypes were scored. Callus images were acquired on a Zeiss Axio Zoom.V16 stereo microscope, and projected areas were quantified in ImageJ.

For *Nicotiana tabacum* shoot induction, fully expanded healthy leaves of *Nicotiana tabacum* were surface sterilized in 75% ethanol for 5 min, rinsed three times with sterile water, incubated in 10% bleach for 7 min, and rinsed five times with sterile water. Sterile leaf discs were punched with a cork borer, blotted dry, and placed abaxial side down on regeneration medium (MS+ 3% sucrose, 1 mg/L 6-BA, 0.1 mg/L NAA, 0.375% Phytagel; pH 5.8). Plates were incubated at 28 °C under a 16-h light/8-h dark photoperiod for ∼4 weeks, with subculture to fresh medium every 14 days. Fo LQFM215 treatments, 10 µL of 10 µM LQFM215 was applied evenly around the perimeter of each disc at day 0. Shoot number per disc was scored at day 30.

Callus induction in *Oryza sativa* ssp. *japonica* (cv. Nipponbare) was performed as previously described ^56^ with minor adaptations. Dehulled mature seeds were surface-sterilized in 50% Clorox solution containing 5 µL Tween-20 for 30 min, rinsed three times with sterile water, and placed on callus-induction 2N6 medium supplemented with 2.0 mg/L 2,4-D. Plates were incubated at 28 °C in darkness for 11 days. Scutellum-derived calli were imaged by Zeiss Axio Zoom.V16 Stereo Microscope and callus area per seed was quantified in ImageJ. Where indicated, 10 µL drop of 10 µM LQFM215 was pipetted directly onto the seed surface immediately after plating.

### Molecular docking and structure prediction

Complexes of Arabidopsis ProT2 (Q9P2962) and ProT3 (Q9SJP9) with L-proline or LQFM215 (ProT2–L-proline, ProT2–LQFM215, ProT3–L-proline, and ProT3–LQFM215) were predicted using AlphaFold v3.0.1^83^ on an NVIDIA A100 GPU. For each protein–ligand complex, five independent models were generated using different random seeds. Protein–ligand contact residues of ProT2 and ProT3 with L-proline or LQFM215 were then analyzed in these models using ChimeraX v1.10.1^84^, with a standard distance cutoff of 4 Å. Colors for different models were selected using the qualitative palette from ColorBrewer 2.0.

## Acknowledgements

We thank L. Xu (CAS Center for Excellence in Molecular Plant Sciences, China), C. H. Khang (University of Georgia, US) for sharing materials. This project is supported by the National Science Foundation (IOS-2039313 to L.Y.) and the National Institutes of Health (R35GM143067 to L.Y.). Support was also provided by the Serrapilheira Institute (G-1811-25705 to P.J.P.L.T.), by the National Council for Scientific and Technological Development (CNPq; 308349/2022-9 to P.J.P.L.T.), and by the São Paulo Research Foundation (FAPESP; 2018/24432-0 to P.J.P.L.T.; 2023/10167-1 to N.C.F.D.). Computational resources were provided in part by the Georgia Advanced Computing Resource Center, a partnership between the University of Georgia’s Office of the Vice President for Research and Office of the Vice President for Information Technology.

## Author contributions

L.Y. and D.X. conceptualized the study. D.X., Z.M.B., A.Z., S.C., Y.Z., and F.K. performed experiments. N.C.F.D., C.J.N., L.W., and M.T.B performed data analyses. D.X., L.Y., D.Y.X., H.H.N.E.A, P.J.P.L.T. and Y.Y. contributed to data interpretation. D.X. and L.Y. wrote the manuscript with input from all authors. All authors reviewed and approved the final manuscript.

---

**Extended Data Fig. 1.**
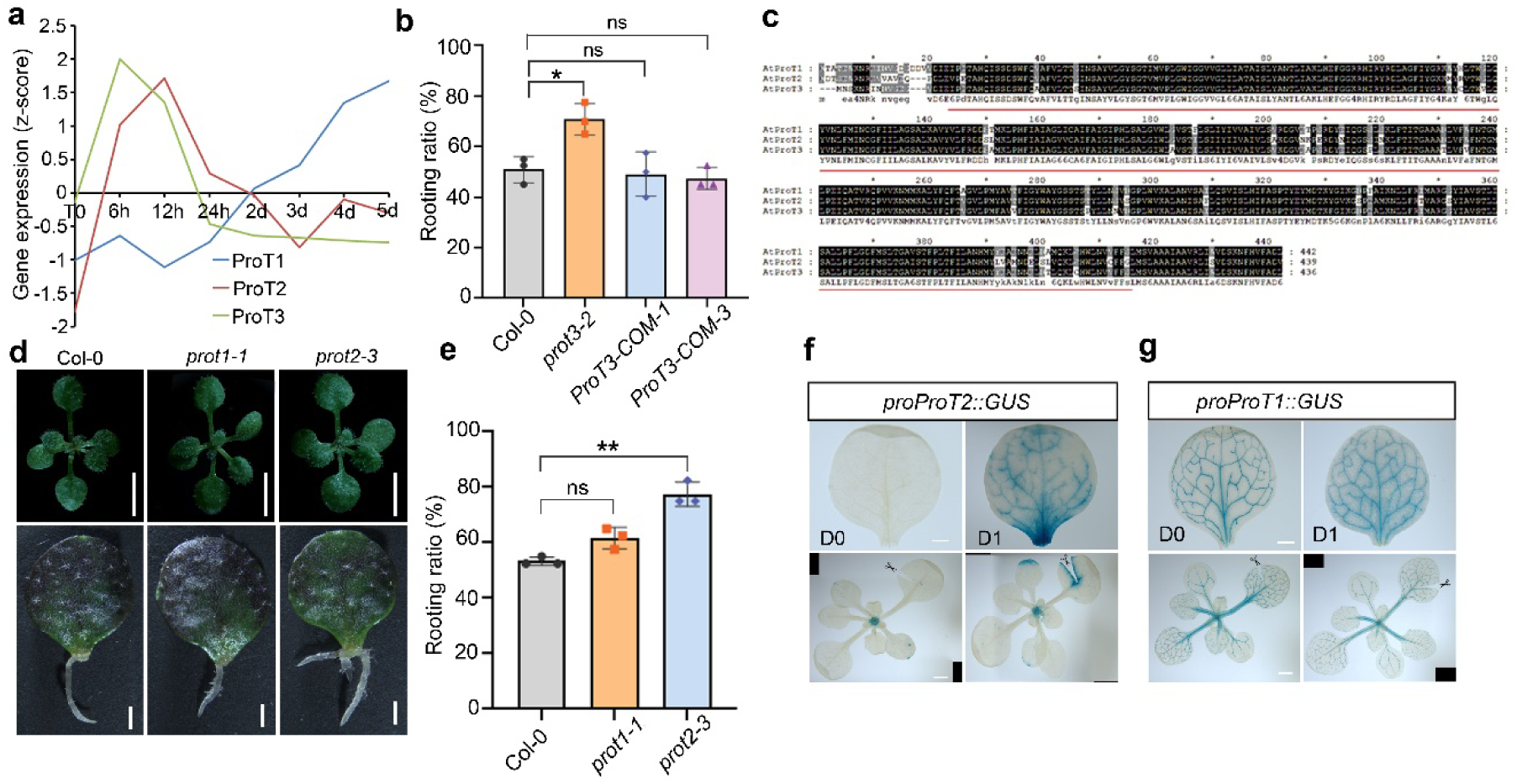
Expression and functional analysis of *ProT* genes. a, Wound-induced expression of *ProT1*, *ProT2* and *ProT3 based on* public transcriptomic datasets ^50^. b, Rooting ratio of Col-0, *prot3-2* and complementation lines (n = 3 biological replicates, 40 explants each). c, Sequence alignment of Arabidopsis *ProTs* and homologs. Red lines indicate transmembrane domain. d, Representative DNRR phenotype of Col-0, *prot1-1* and *prot2-3* after 10 DAC; Scale bars, 0.5 cm (top) and 1 mm (bottom). e, Rooting ratio of Col-0, *prot1-1* and *prot2-3* (n = 3 biological replicates, 40 explants each). f, g, GUS staining of *proProT2::GUS* (f) and *proProT1::GUS* (g) at D0 and D1 cultured on B5 medium. scale bars, 500 µm (top) and 1 mm (bottom). Data presented as mean ± s.d.; *n* denotes the number of biological replicates. Statistical significance was determined by two-tailed Student’s *t*-test; *P < 0.05, **P <0.01, ***P < 0.001; ns, not significant.

**Extended Data Fig. 2.**
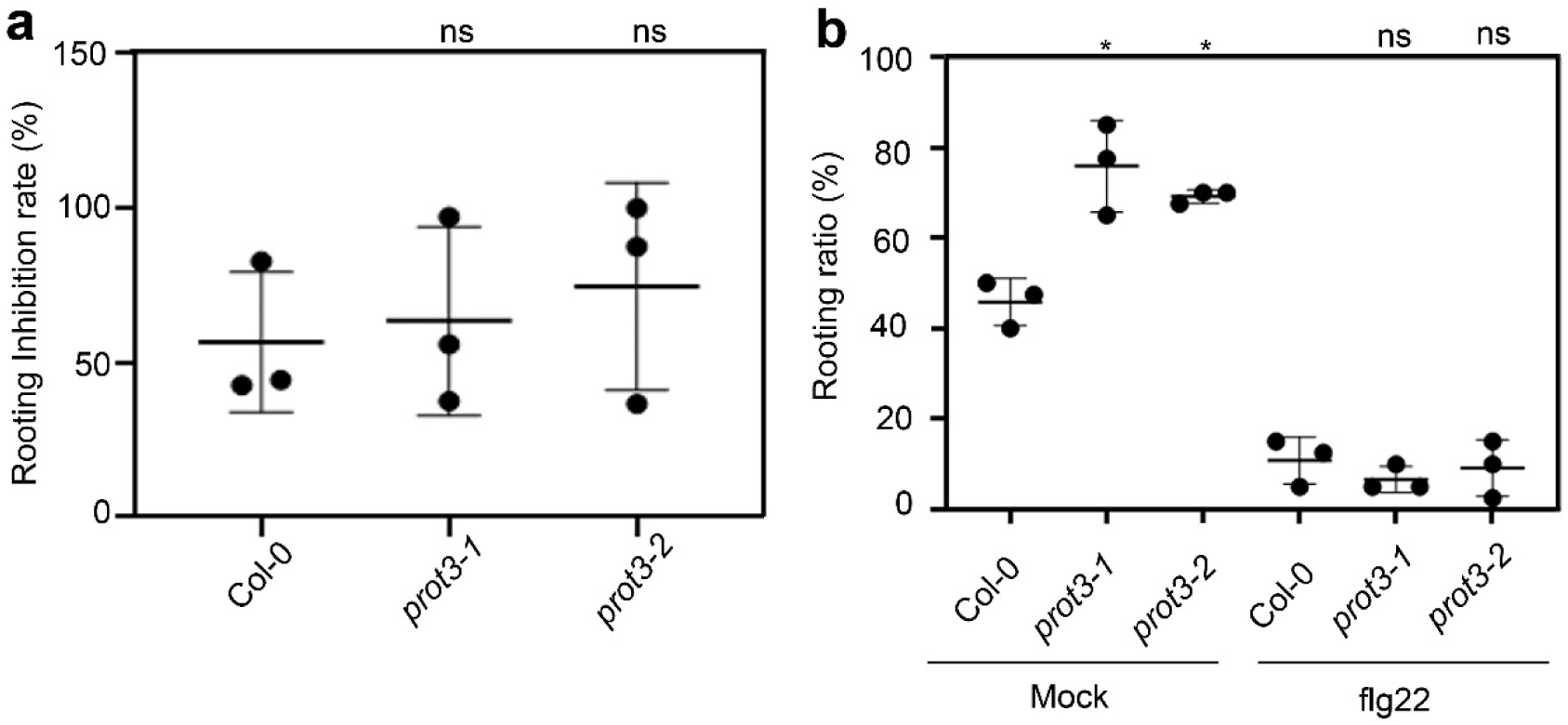
Rooting response of *prot3* mutants under biotic stress conditions. a, Rooting inhibition rate of Col-0, *prot3-1* and *prot3-2* leaf explants upon *hrrC^-^* treatment. (n = 3 biological replicates, 40 explants each). hrcC^-^ was applied at a concentration of OD600=0.01. b, Rooting ratio of Col-0, *prot3-1* and *prot3-2* leaf explants cultured on mock or flg22-supplemented medium for 10 days. (n = 3 biological replicates, 40 explants each). Data presented as mean ± s.d.; *n* denotes the number of biological replicates. Statistical significance was determined by two-tailed Student’s *t*-test; *P < 0.05, **P <0.01, ***P < 0.001; ns, not significant.

**Extended Data Fig. 3.**
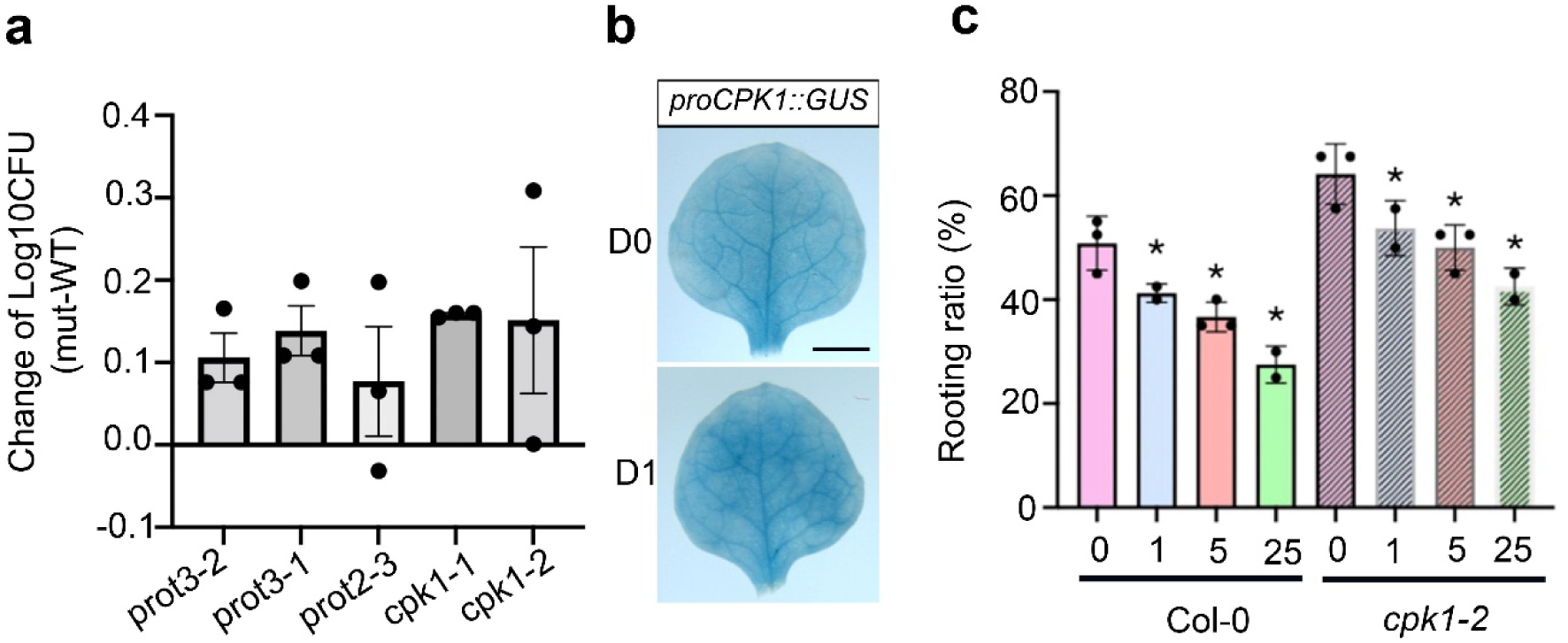
CPK1 function in DNRR and defense. a, Summary of bacteria growth curve from multiple experiments. Each dot represents the log_10_CFU difference between Col-0 and mutant in one independent experiment. *prot3*, *prot2* and *cpk1* mutants showed mild enhance of susceptibility to *Pto* DC3000. b, *proCPK1::GUS* expression is not induced by wounding. GUS staining of *proCPK1::GUS* Arabidopsis leaves at day 0 (D0, a) and 1 day after cutting (D1, b) cultured on B5 medium. No detectable induction of *CPK1* promoter activity was observed following wounding. Scale bar, 1 mm. c, Rooting ratio of Col-0 and *cpk1-2* in response to increasing concentration of NaSA.

**Extended Data Fig. 4.**
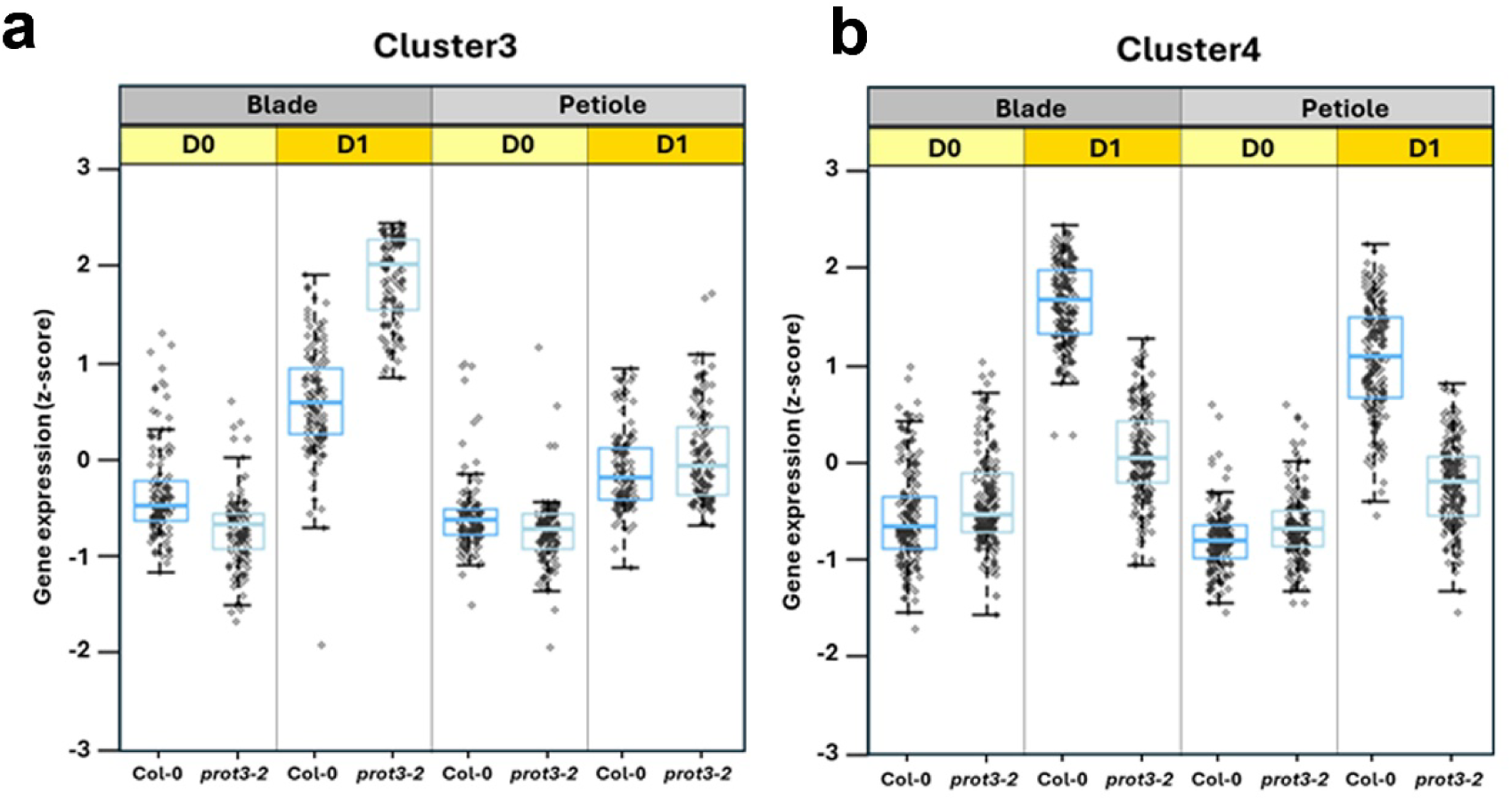
Expression patterns of gene clusters affected by ProT3 in Arabidopsis leaf explants. a, Genes in cluster 3 were more strongly induced in the leaf blades of prot3-2 at 1 day after cutting (D1). b, Genes in cluster 4 displayed lower activation levels in both blades and petioles of *prot3-2* compared with Col-0 at D1.

**Extended Data Fig. 5.**
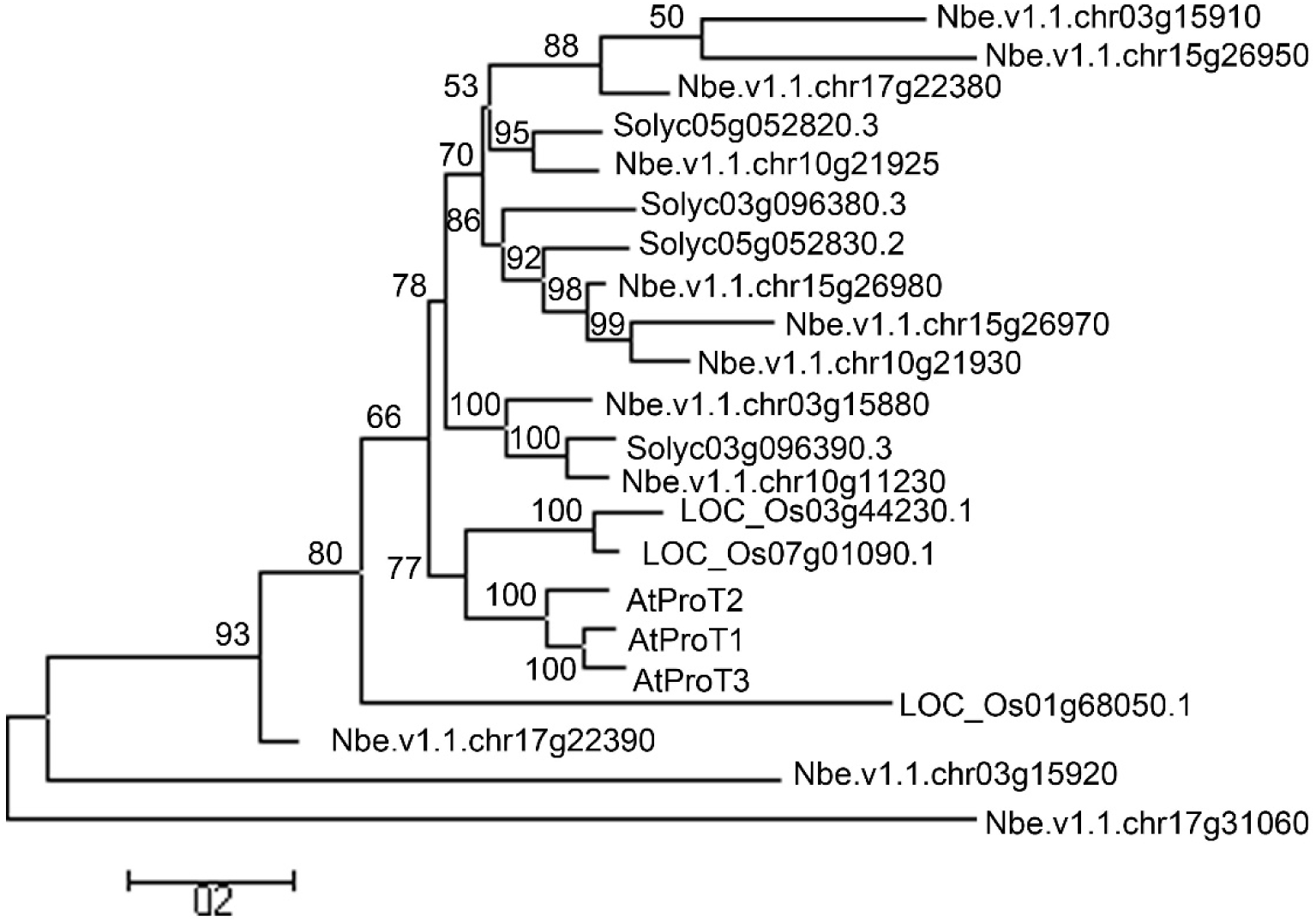
Phylogenetic analysis of proline transporter (ProT) homologs from Arabidopsis, tomato, rice, and *Nicotiana benthamiana*. A neighbor-joining phylogenetic tree showing the relationships among predicted proline transporter (ProT) homologs from *Arabidopsis thaliana* (AtProT1–3), *Solanum lycopersicum* (Solyc), *Oryza sativa* (LOC_Os), and *Nicotiana benthamiana* (Nbe). The tree was constructed using MEGA 3.1 with 1,000 bootstrap replicates, and bootstrap support values are indicated at each node. The scale bar represents 0.2 substitutions per site.

**Extended Data Fig. 6.**
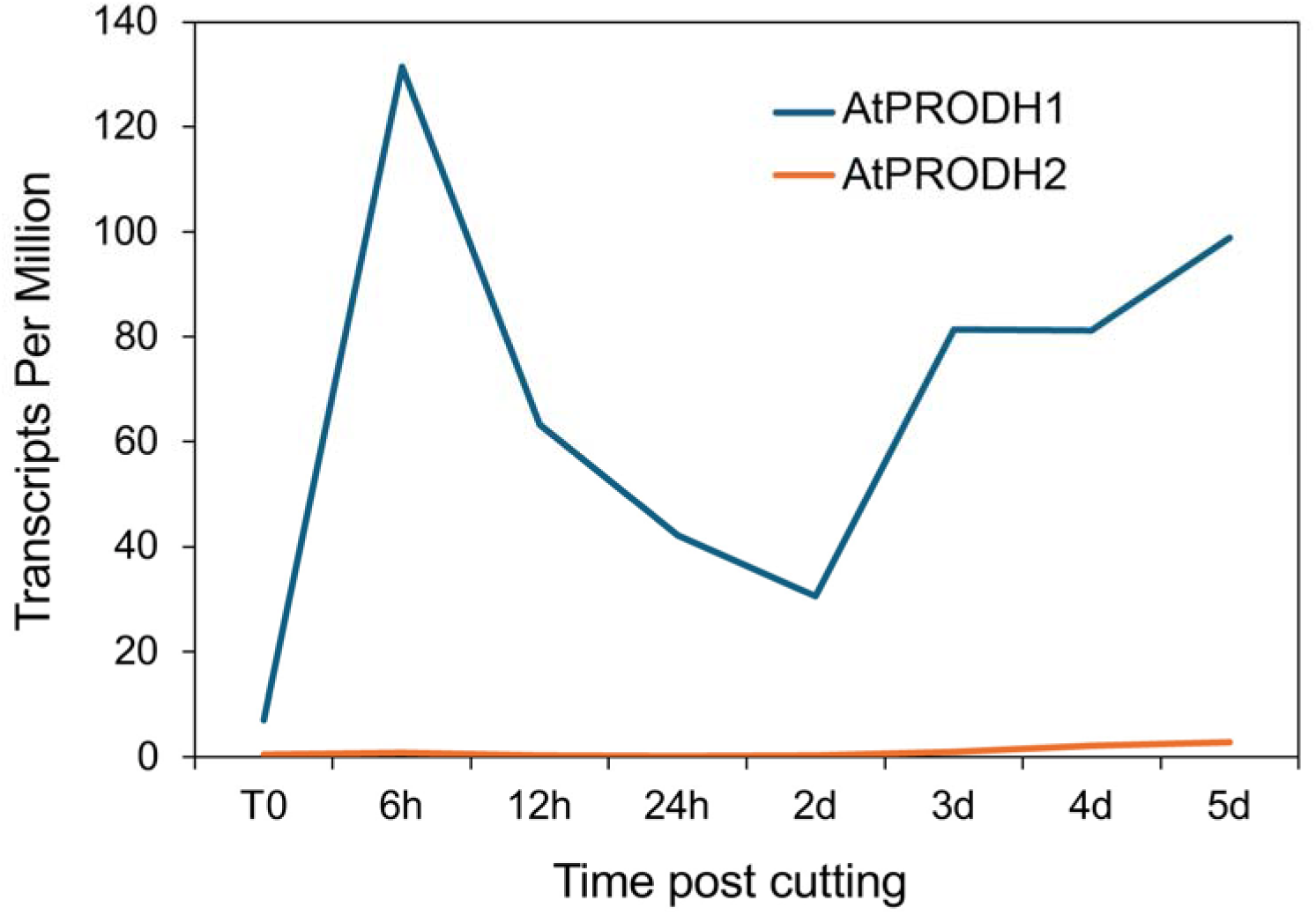
Wound-induced expression of AtProDH1 (AT3G30775) and AtProDH2 (AT5G38710) based on public transcriptomic datasets. Temporal expression patterns of AtPRODH1 and AtPRODH2 following leaf cutting. Transcript abundance (transcripts per million, TPM) was extracted from the public wound-induced regeneration RNA-seq dataset ^50^. AtPRODH1 shows a rapid induction within 6 h after wounding, followed by a gradual decline, whereas AtPRODH2 remains at a low basal level throughout the time course.

